# Genetic Variants and Functional Pathways Associated with Resilience to Alzheimer’s Disease

**DOI:** 10.1101/2020.02.19.954651

**Authors:** Logan Dumitrescu, Emily R. Mahoney, Shubhabrata Mukherjee, Michael L. Lee, William S. Bush, Corinne D. Engelman, Qiongshi Lu, David W. Fardo, Emily H. Trittschuh, Jesse Mez, Catherine Kaczorowski, Hector Hernandez Saucedo, Keith F. Widaman, Rachel Buckley, Michael Properzi, Elizabeth Mormino, Hyun-Sik Yang, Tessa Harrison, Trey Hedden, Kwangsik Nho, Shea J. Andrews, Doug Tommet, Niran Hadad, R. Elizabeth Sanders, Douglas M. Ruderfer, Katherine A. Gifford, Annah M. Moore, Francis Cambronero, Xiaoyuan Zhong, Neha S. Raghavan, Badri Vardarajan, The Alzheimer’s Disease Neuroimaging Initiative (ADNI), Alzheimer’s Disease Genetics Consortium (ADGC), A4 Study Team, Margaret A. Pericak-Vance, Lindsay A. Farrer, Li-San Wang, Carlos Cruchaga, Gerard Schellenberg, Nancy J. Cox, Jonathan L. Haines, C. Dirk Keene, Andrew J. Saykin, Eric B. Larson, Reisa A. Sperling, Richard Mayeux, David A. Bennett, Julie A. Schneider, Paul K. Crane, Angela L. Jefferson, Timothy J. Hohman

**Author notes:** Address Correspondence to: Timothy J Hohman, PhD, Vanderbilt Memory & Alzheimer’s Center, Vanderbilt University Medical Center, 1207 17^th^ Ave S, Nashville, TN 37212. Data used in preparation of this article were obtained from the Alzheimer’s Disease Neuroimaging Initiative (ADNI) database (adni.loni.usc.edu). As such, the investigators within the ADNI contributed to the design and implementation of ADNI and/or provided data but did not participate in analysis or writing of this report. A complete listing of ADNI investigators can be found at: http://adni.loni.usc.edu/wp-content/uploads/how_to_apply/ADNI_Acknowledgement_List.pdf.

## Abstract

Approximately 30% of older adults exhibit the neuropathologic features of Alzheimer’s disease (AD) without signs of cognitive impairment. Yet, little is known about the genetic factors that allow these potentially resilient individuals to remain cognitively normal in the face of substantial neuropathology. We performed a large, genome-wide association study (GWAS) of two previously validated metrics of cognitive resilience quantified using a latent variable modeling approach and representing better-than-predicted cognitive performance for a given level of neuropathology. Data were harmonized across 5,108 participants from a clinical trial of AD and three longitudinal cohort studies of cognitive aging. All analyses were run across all participants and repeated restricting the sample to individuals with normal cognition to identify variants at the earliest stages of disease. As expected, all resilience metrics were genetically correlated with cognitive performance and education attainment traits (p-values<2.5×10^−20^), and we observed novel correlations with neuropsychiatric conditions (p-values<7.9×10^−4^). Notably, neither resilience metric was genetically correlated with clinical AD (p-values>0.42) nor associated with *APOE* (p-values>0.13). In single variant analyses, we observed a genome-wide significant locus among participants with normal cognition on chromosome 18 upstream of *ATP8B1* (index SNP rs2571244, MAF=0.08, p=2.3×10^−8^). The top variant at this locus (rs2571244) was significantly associated with methylation in prefrontal cortex tissue at multiple CpG sites, including one just upstream of *ATPB81* (cg19596477; p=2×10^−13^). Overall, this comprehensive genetic analysis of resilience implicates a putative role of vascular risk, metabolism, and mental health in protection from the cognitive consequences of neuropathology, while also providing evidence for a novel resilience gene along the bile acid metabolism pathway.

Furthermore, the genetic architecture of resilience appears to be distinct from that of clinical AD, suggesting that a shift in focus to molecular contributors to resilience may identify novel pathways for therapeutic targets.

## Introduction

Alzheimer’s disease (AD) is characterized by the presence of neuritic plaques and neurofibrillary tangles in the brain at autopsy. Clinically, it presents with progressive cognitive impairment. Yet, due to the long prodromal period of AD and unknown biological factors, not everyone with AD neuropathology presents with cognitive impairment. In fact, among cognitively normal volunteers agreeing to autopsy at the time of death, 70% have varying degrees of AD pathology (Sonnen *et al.*, 2011), and 30% have sufficient neuropathology in their brain to meet neuropathological criteria for AD (i.e., “Asymptomatic AD”) (Rahimi and Kovacs, 2014). Identifying the molecular factors that underlie the resilience observed in asymptomatic AD may provide novel therapeutic targets for clinical intervention and provide additional insight into the genetic architecture of AD.

While there has been some prior discovery work using genomic data (Mostafavi *et al.*, 2018; Yu *et al.*, 2018), previous work characterizing the genetic contributors to asymptomatic AD has primarily focused on candidate genes (Monsell *et al.*, 2013; Monsell *et al.*, 2017; Franzmeier *et al.*, 2019) due to the lack of sufficient sample size to complete full genome-wide analyses. A major barrier in moving analyses forward has been the categorical definitions of asymptomatic AD that drastically reduce the number of participants available for analysis. In the last decade, residual approaches to quantifying continuous metrics of “resilience” have emerged as potential endophenotypes for genetic analyses (Yu *et al.*, 2015; White *et al.*, 2017; Boyle *et al.*, 2019). The basic approach is to deconvolve cognitive scores into components that are explained and unexplained by proxy or direct measures of neuropathology (Reed *et al.*, 2010). These residual approaches model better-than and worse-than predicted cognitive performance to represent higher vs. lower resilience (Yu *et al.*, 2015; Boyle *et al.*, 2019). Recently, our group has extended these residual approaches to quantify and validate continuous metrics of “cognitive resilience” (representing better-than-predicted cognitive performance given an individual’s burden of AD neuropathology) and “brain resilience” (representing better-than-predicted brain volumes given an individual’s burden of AD neuropathology) (Hohman *et al.*, 2016b). These continuous metrics are strong predictors of future cognitive decline and cognitive impairment (Hohman *et al.*, 2016b). The goal of the present analysis was to evaluate genetic predictors of cognitive resilience across the genome.

A few genome-wide analyses have been completed that focus on resilience in asymptomatic AD, although with limited sample sizes (Hohman *et al.*, 2014a; Hohman *et al.*, 2014b; Hohman *et al.*, 2016a; White *et al.*, 2017). Recently, approximately 3,000 samples with both whole-genome genetic data and *in vivo* brain measures of amyloid burden from the Anti-Amyloid Treatment in Asymptomatic AD (A4) clinical trial were made publicly available, providing an unmatched resource for exploring the genetics of resilience to AD. We performed the largest (N=5,108) genome-wide association study (GWAS) of cognitive resilience in AD by leveraging harmonized resilience metrics across the cross-sectional A4 study and three longitudinal cohort studies of AD. Validation of identified genomic candidates was completed using gene expression data from post mortem brain tissue and genotype data from large-scale case/control datasets of AD. Importantly, we also performed comprehensive genetic correlation and pathway analyses to provide critical information about the fundamental biological pathways that may protect the brain from the downstream consequences of AD neuropathology.

## Materials and Methods

### Participants

Participant data was acquired from multiple cohort studies including screening data from the Anti-Amyloid Treatment in Asymptomatic Alzheimer’s Disease (A4) Study, the Alzheimer’s Disease Neuroimaging Initiative (ADNI), the Religious Orders Study and Rush Memory & Aging Project (ROS/MAP), and the Adult Changes in Thought (ACT) Study. The <u>A4</u> Study screening data were acquired as part of a clinical trial that began in 2014 (Sperling *et al.*, 2014). All participants were recruited with normal cognition, and amyloid Positron Emission Tomography (PET) imaging was performed at screening. Additionally, participants with a Delayed Logical Memory score less than 6 or greater than 18 were excluded from PET scans and are not included in the present analysis. <u>ADNI</u> was launched in 2003 and over the four phases of the study now includes >1,800 individuals age 55-90 (www.adni-info.org). Recruitment was designed to mimic clinical trials and therefore included individuals with normal cognition, mild cognitive impairment, and AD at baseline. Data from ADNI-1, ADNI-2, and ADNI-GO are included in the present analyses. <u>ACT</u> began in 1994 and recruited a random sample of nondemented older adults from the Seattle metropolitan area (Kukull *et al.*, 2002). A subset of participants in ACT agreed to brain donation and are included in these analyses. <u>ROS</u> launched in 1994 and recruited Catholic nuns, priests, and brothers from across the United States, and <u>MAP</u> launched in 1997 and recruited cognitively normal older adults from the Chicago metropolitan area (Bennett *et al.*, 2018). Those who agreed to brain donation are included in the present analysis.

### Amyloid PET Acquisition

For ADNI and A4 participants, amyloid burden was quantified using amyloid PET. PET procedures in ADNI are described at the ADNI website (http://www.adni-info.org). A4 and ADNI are both largescale multi-site studies for which PET amyloid acquisition was completed on multiple platforms, including GE, Philips, and Siemens. In all cases, PET data were acquired using a dynamic 3D scan with four 5-minute frames acquired 50-70 minutes post injection. A subset of ADNI participant data were acquired using ^11^C-PiB, but the majority of ADNI and all of A4 was acquired using ^18^F-Florbetapir. Standardized uptake value ratios (SUVR) were quantified relative to whole cerebellum, and a composite mean SUVR was quantified across cortical regions as a summary metric of amyloid burden.

### Amyloid PET Processing and Harmonization

Harmonization of amyloid PET levels was performed using composite cortical values calculated within ADNI and A4 separately. We applied a Gaussian Mixture Model (GMM) within each dataset to place values on the same scale using a recently developed harmonization algorithm (Properzi *et al.*, 2019). GMMs were estimated among cognitively normal individuals using a two-component model fit and applied to the entire sample. Mean SUVRs were scaled and normalized using the mean and standard deviation estimated from the predicted amyloid negative gaussian distribution. A more comprehensive assessment of this and alternative harmonization approaches was recently published by our group (Raghavan *et al.*, 2020), but we used the present approach because it makes the fewest assumptions about the data and was more robust to outliers than alternative approaches. The final scaled score represents a z-score based on the predicted amyloid negative distribution among cognitively normal older adults.

### Postmortem Assessment of Neuropathology

For ACT and ROS/MAP participants, neuritic plaque burden was quantified with CERAD scores. A comprehensive neuropathological evaluation was completed at each site, including full CERAD staging as previously described (Mirra *et al.*, 1991).

### Neuropsychological Composites

Harmonization of cognitive tests in ADNI and A4 was completed using the Preclinical Alzheimer Cognitive Composite (PACC), calculated in each dataset individually using item level data from Logical Memory Immediate and Delayed Recall, WAIS-R Digit Symbol Substitution Test, the Mini-Mental State Exam, and the Selective Reminding Test or the delayed word recall from the ADAS-Cog. In all four datasets, a previously published protocol to harmonize neuropsychological scores in the domains of memory and executive function was used (Crane *et al.*, 2017). A memory composite was calculated in all four datasets, and an executive function composite was quantified in ACT, ADNI, and ROS/MAP (there was insufficient item level data in A4). A detailed description of the item level data and model that was included in these composite metrics is presented in **Supplementary Methods**.

### Quantification of Resilience Metrics

Resilience metrics were quantified using established procedures (Hohman *et al.*, 2016b) and the model is presented in **Fig. 1**. Briefly, individual regression models estimated amyloid pathology associations with cognition covarying for age and sex. A robust weighted least squares estimator in a confirmatory factor analysis was quantified using Mplus (Muthén and Muthén, 1998-2015) (version 7.31) to summarize residuals from the linear regression models into composite measures representing the degree to which an individual performed better or worse than predicted given their age, sex, and amyloid load (note that years of education was integrated into the second order latent trait). The outcomes of interest were Residual Cognitive Resilience and Combined Resilience where Residual Cognitive Resilience was quantified from residuals and Combined Resilience was summarized as the covariance of educational attainment with Residual Cognitive Resilience. A detailed description of the methodology and quantified resilience metrics is presented in **Supplementary Methods**.

**Figure 1.**
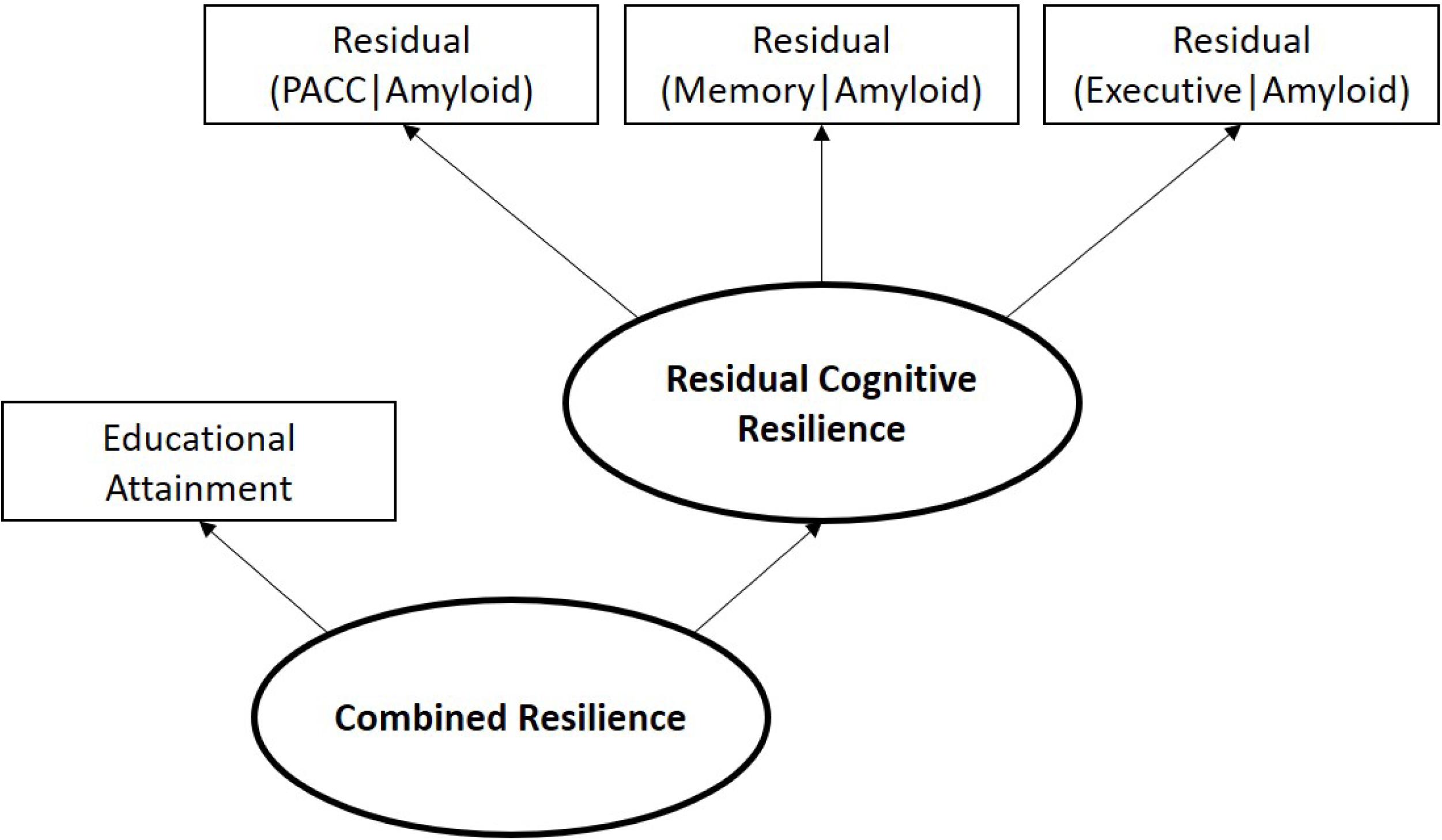
Quantification of Resilience Metrics. Residuals from linear regression models in which a cognitive score was regressed on age, sex, and amyloid levels were extracted and entered as indicator variables in a partial least squares path model using established procedures. Combined Resilience was quantified as a second order latent trait in the model in which educational attainment was included as an additional indicator variable.

### Genotype Processing and Quality Control

Genotyping in all cohorts was performed using DNA extracted from whole blood or brain tissue on different genotyping arrays. For A4, the Illumina Global Screening Array was used for genotyping. ACT participants were genotyped on an Illumina Human660W-Quad. Three Illumina platforms were used in ADNI: Human610-Quad, HumanOmniExpress, and Omni 2.5M. ROSMAP genotypes were also obtained on three platforms: Affymetrix Genechip 6.0, Illumina Human1M, and Illumina Global Screening Array. In ADNI and ROSMAP, sample sets genotyped on different arrays were processed and imputed in parallel and merged after imputation. Quality control (QC) was performed using standard procedures, including removal of SNPs and samples with >5% genotype missingness, removal of SNPs with <1% minor allele frequency (MAF) or Hardy-Weinberg Equilibrium (HWE) p-values <10^−6^, and removal of samples with sex discrepancies, cryptic relatedness (pi-hat >0.25), or who were not non-Hispanic White by self-report or by population principal component (PC) analysis.

Genotypes were then imputed with Minimac3 on the Michigan Imputation Server (https://imputationserver.sph.umich.edu) using the HRC r1.1 2016 reference panel. Post-imputation QC steps included removal of SNPs with imputation quality score R^2^ <0.90, call rate <95%, MAF <1%, or HWE p-value <10^−6^. Imputed datasets were then merged for the two autopsy cohorts (ACT and ROS/MAP) and the two PET imaging cohorts (A4 and ADNI). Non-overlapping SNPs (i.e., those with missingness >95%) were excluded. A total of 4,840,740 SNPs remained and were included in the analysis.

### Statistical Analyses

Our analysis workflow is presented in **Fig. 2**. Following phenotype harmonization and calculation of resilience metrics (i.e., Residual Cognitive Resilience and Combined Resilience) for each cohort, genome-wide association analyses were completed using linear regression in PLINK (version 1.9, https://www.cog-genomics.org/plink/1.9). GWAS was performed in the combined autopsy dataset and the combined PET dataset. For each dataset, two models were run. The first model estimated resilience among individuals across the spectrum of dementia, including individuals with normal cognition, mild cognitive impairment, and AD. The second model restricted the sample to individuals with normal cognition to focus on resilience during the preclinical phase of disease. In all models, covariates included age, sex, and the first three population PCs. The genome-wide threshold for statistical significance was set *a priori* at α=5×10^−8^. Summary statistics at each marker across the autopsy dataset and the PET dataset were then combined in a fixed-effect meta-analysis using the GWAMA software program (Mägi and Morris, 2010).

**Figure 2.**
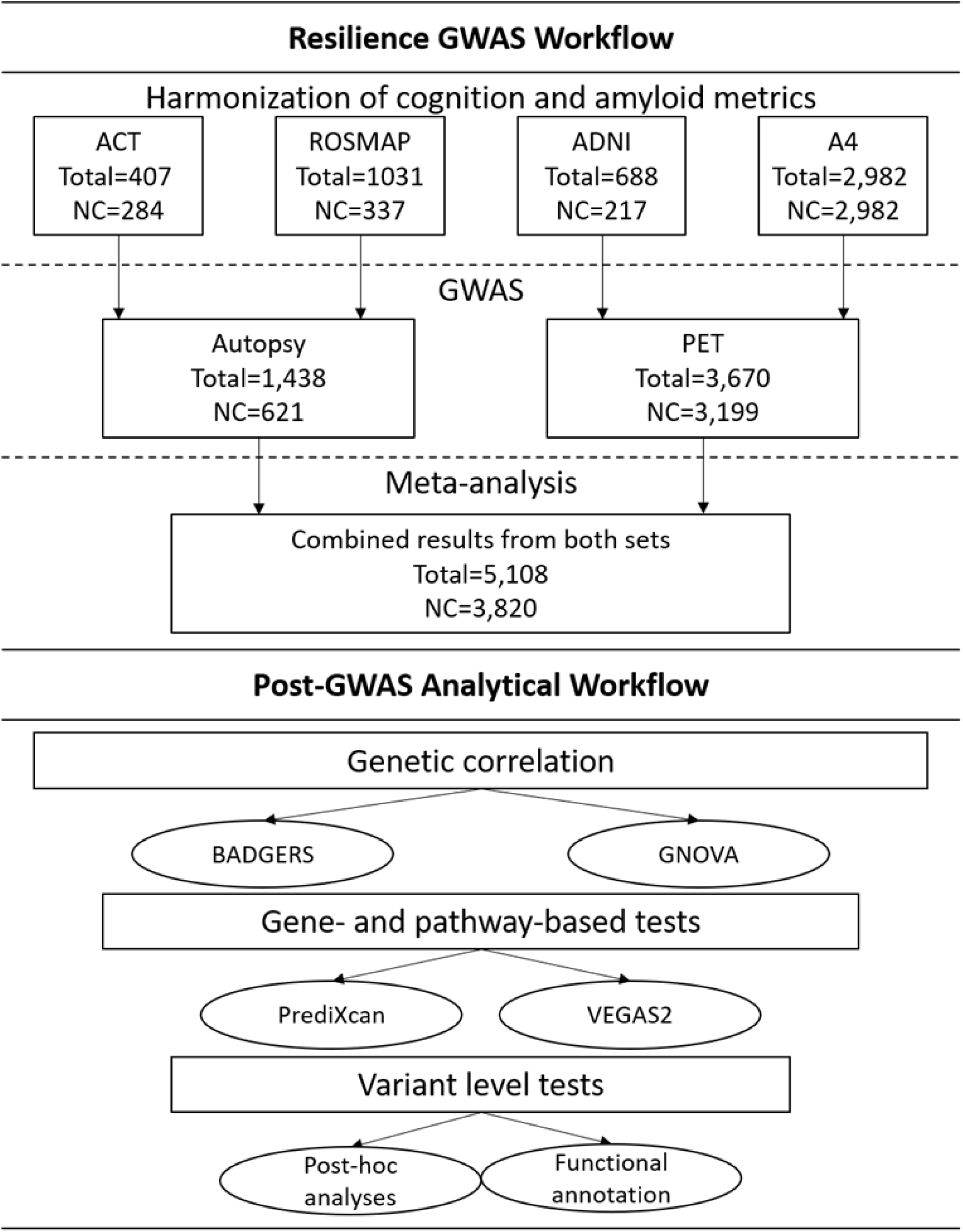
Workflow of Analytical Activities.

We first summarized genetic signal across the genome using summary statistics from our resilience GWAS to estimate genetic correlations between resilience phenotypes and 67 complex traits with publicly accessible GWAS summary statistics using the Genetic Covariance Analyzer (GNOVA) program (Lu *et al.*, 2017). Details about the source of summary statistics for each trait are presented in **Supplementary Table 1**. This provided a first level of validation that the genetic signal in our analysis was correlated with common phenotypes (e.g., cognitive performance and educational attainment) while also providing insight into novel resilience associations. Additionally, we replicated our top genomic correlation results leveraging the BADGERS program (Yan *et al.*, 2018) and quantified correlation across 1,738 traits in the UK Biobank (http://biobank.ndph.ox.ac.uk/showcase/). To aid in interpretation of genetic covariance results, we also quantified heritability estimates using the Genome-wide Complex Trait Analysis (GCTA) tool (Yang *et al.*, 2011). Heritability of each resilience phenotype was quantified within the PET and Autopsy datasets separately, and within a combined dataset including all samples. Estimates were quantified across all participants and when restricting the sample to individuals with normal cognition.

**Table 1.**
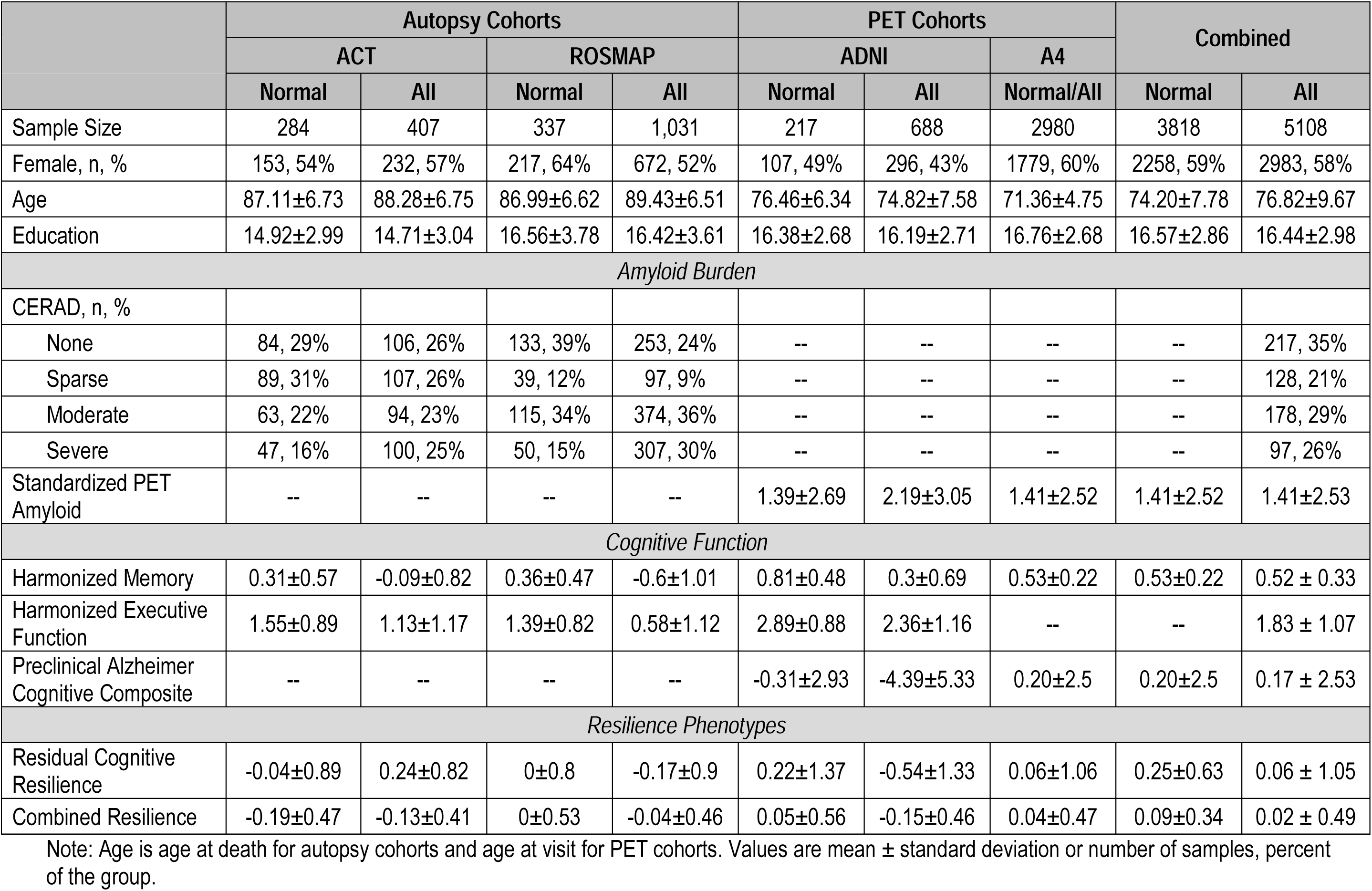
Participant Characteristics.

Next, we performed gene- and pathway-level analyses using VEGAS2 (Liu *et al.*, 2010; Mishra and Macgregor, 2015; Mishra and MacGregor, 2017) and PrediXcan (Gamazon *et al.*, 2019). PrediXcan models were estimated for 44 tissues in the GTEx Portal and for additional disease relevant tissues, including prefrontal cortex from CommonMind and monocytes from the Multi-Ethnic Study of Atherosclerosis (MESA). Correction for multiple comparisons in gene-level analyses was quantified using the false discovery rate (FDR) procedure, which accounted for all 258,562 gene-tissue combinations. The *a priori* threshold for significance of the VEGAS pathway results was p<1×10^−5^, which was based on a simulation-derived 95% empirical significance threshold taking into account the multiple testing of 6,213 correlated pathways (Mishra and MacGregor, 2017).

Finally, single variant GWAS loci were mapped to genes and functionally annotated leveraging INFERNO (http://inferno.lisanwanglab.org/) (Amlie-Wolf *et al.*, 2018) and the Brain xQTL Serve database (http://mostafavilab.stat.ubc.ca/xqtl/) (Ng *et al.*, 2017). INFERNO integrates hundreds of publicly available functional genomics databases, including databases of transcription factor binding sites, expression quantitative trail loci (eQTL), and enhancer activity. The Brain xQTL Serve database includes additional eQTL, methylation-QTL (mQTL), and histone-QTL (hQTL) analyses.

## Results

In total, 5,108 individuals across the four cohorts (A4 n=2,982; ROS/MAP n=1,031; ADNI n=688; ACT n=407) had both genome-wide genotype and resilience phenotype data, 3,820 (75%) of whom were cognitively normal. Participant characteristics are presented in **Table 1**. In general, participants were mostly female (with the exception of ADNI) and were well-educated. Individuals in the PET cohorts tended to be younger than individuals in the autopsy cohorts.

### Genetic Covariance Results

Heritability estimates for each resilience phenotype are presented in **Supplementary Table 2.** Briefly, we observed larger heritability estimates when restricting the sample to individuals with normal cognition (Residual Cognitive Resilience h^2^=0.20-0.28, Combined Resilience h^2^=0.23-0.99) compared to the entire sample (Residual Cognitive Resilience h^2^=0.00-0.08, Combined Resilience h^2^=0.19-0.67).

Using the summary statistics from the resilience GWAS, we performed genetic covariance analyses to gain insight into any shared genetic basis of relevant biological processes. Pair-wise genetic covariances between Combined Resilience GWAS results in all participants and 67 health-related phenotypes are depicted in **Fig 3** and presented in **Supplementary Table 3**. Ten genetic correlation analyses survived correction for multiple testing. We observed strong and expected positive correlations with cognitive performance and educational attainment (p<1.4×10^−19^), validating our metric and providing strong evidence of consistency in the observed polygenic signal across comparable measures from independent datasets.

**Figure 3.**
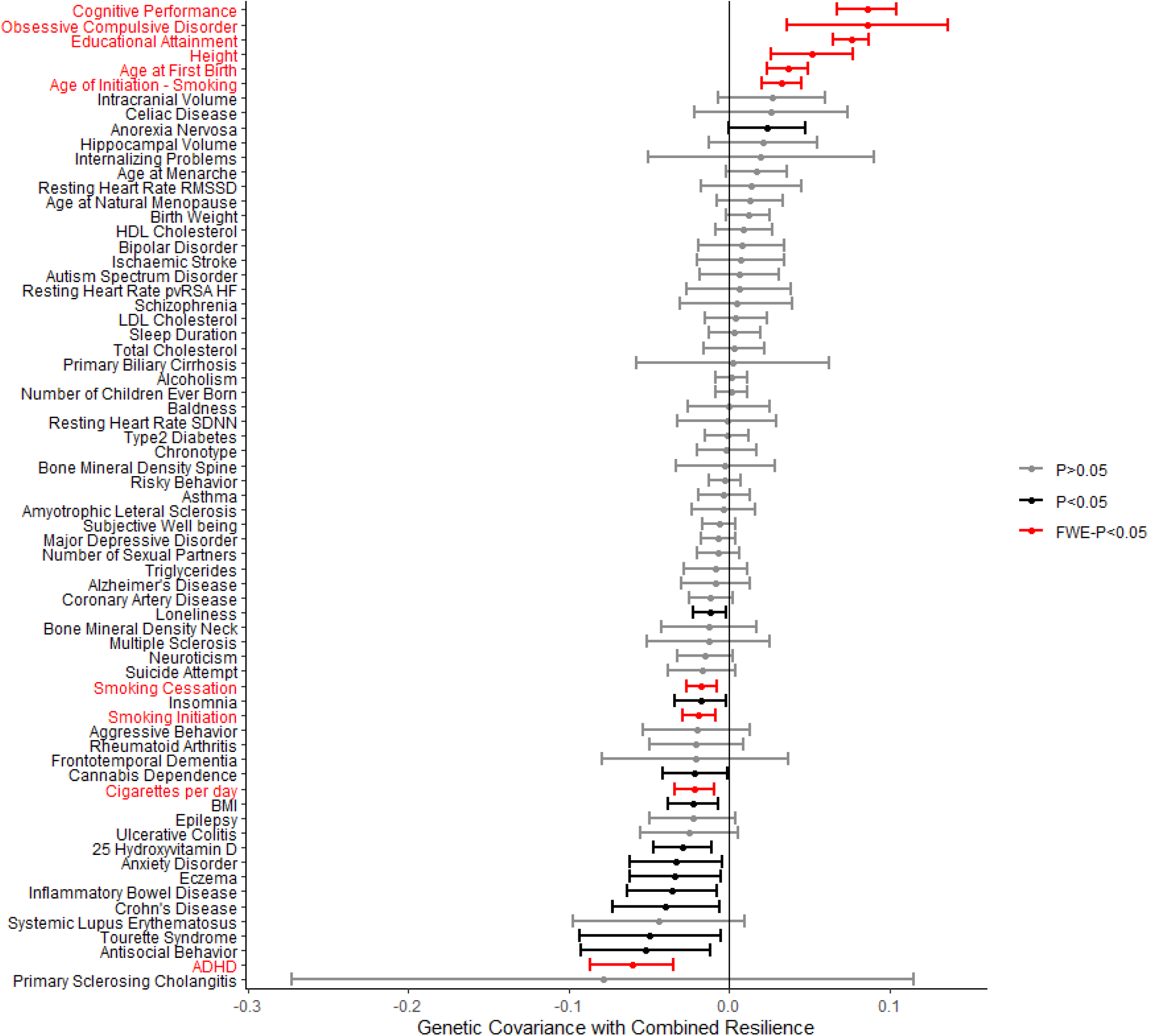
Genome-Wide Genetic Covariance Results. Genetic covariances between Combined Resilience and 67 complex traits. Error bars represent 95% confidence intervals. FWE-P: corrected p-value based on the family-wise error rate.

Additionally, we observed multiple novel correlations, including two smoking behavior phenotypes: age at smoking initiation (genetic correlation=0.033; p=2.0×10^−7^) and number of cigarettes per day (genetic correlation=-0.021; p=8.0×10^−4^). Additional novel correlations included two neuropsychiatric conditions, whereby increased genetic risk of obsessive compulsive disorder (OCD) was correlated with higher levels of resilience (p=7.9×10^−4^) while increased genetic risk of attention deficit hyperactivity disorder (ADHD) was associated with lower levels of resilience (p=4.7×10^−6^). Interestingly, older age at first birth was associated with higher levels of resilience (p=1.1×10^−8^). Genetic correlations with Residual Cognitive Resilience were very similar to those observed for Combined Resilience and were similar when restricting the sample to cognitively normal individuals (**Supplementary Tables 3-4**).

As a second level of validation, we also quantified genetic correlations with phenotypes in the UK Biobank leveraging a recently published method to perform phenome-wide association analyses leveraging summary statistics (Yan *et al.*, 2018). Consistent with GNOVA results, we observed strong correlations with numerous education and cognitive phenotypes (**Supplementary Tables 5-6**). We also verified correlations with age at first birth (p=6.2×10^−12^) and observed some intriguing novel correlations.

Interestingly, there was no evidence for genetic correlation between resilience phenotypes and clinical AD (p=0.45). Similarly, when evaluating the 40 previously identified AD risk variants from approximately 25 loci (Lambert *et al.*, 2013; Jansen *et al.*, 2019; Kunkle *et al.*, 2019), only three SNPs showed nominal evidence of association with either resilience phenotype (**Supplementary Table 7**). Similar results were also observed when fully analyzing the *APOE* haplotype, whereby increasing numbers of *APOE* ε4 alleles or number of *APOE* ε2 alleles were not associated with either resilience phenotype (p-values>0.13). Together these results suggest the polygenic signal underlying the resilience phenotypes is distinct from clinical AD.

### Gene-Level and Pathway Results

Next, we continued to explore the genetic architecture of resilience on both a gene and pathway level. Gene-level results in individual tissues and cross-tissue, based on predicted gene expression associations with resilience, are reported in **Supplementary Tables 8-11**. Resilience metrics were not associated with predicted gene expression among individual tissues or across tissues after Bonferroni correction for multiple testing. The most significant gene in the cross-tissue analyses was *ZNF451*, which was associated with Combined Resilience in individuals with normal cognition at p<6.6×10^−6^ (**Supplementary Table 9**).

In pathway-level analyses using VEGAS2, no molecular pathways remained significant when correcting for multiple comparisons. However, when restricting to cognitively normal participants in the Combined Resilience analysis, there was nominal evidence of enrichment in the dehydrogenase pathway (p=2.5×10^−5^; PANTHER database) and the amino acid metabolism pathway (8.7×10^−5^; PANTHER database).

### Single-Variant Associations with Resilience

Finally, we focused on single variant level analyses to identify novel genetic loci associated with resilience. Genome-wide significant results are presented in **Fig. 4a**, and detailed results for all models are presented in **Supplementary Tables 12-15**. When including all diagnoses in the GWAS, we did not observe any variants that reached statistical significance in either Residual Cognitive Resilience or Combined Resilience analyses. When restricting analyses to individuals with normal cognition, we identified a locus on chromosome 18 just upstream of the *ATP8B1* gene that reached genome-wide significance in Combined Resilience analyses (**Fig. 4b**). More specifically, the minor allele of the index SNP at this locus (rs2571244; MAF=0.08) was associated with lower levels of Combined Resilience (β=-0.11, p=2.3×10^−8^), and the direction of association was consistent across the PET and Autopsy datasets (**Fig. 4c**). No genome-wide associations were observed in the Residual Cognitive Resilience analyses among participants with normal cognition.

**Figure 4.**
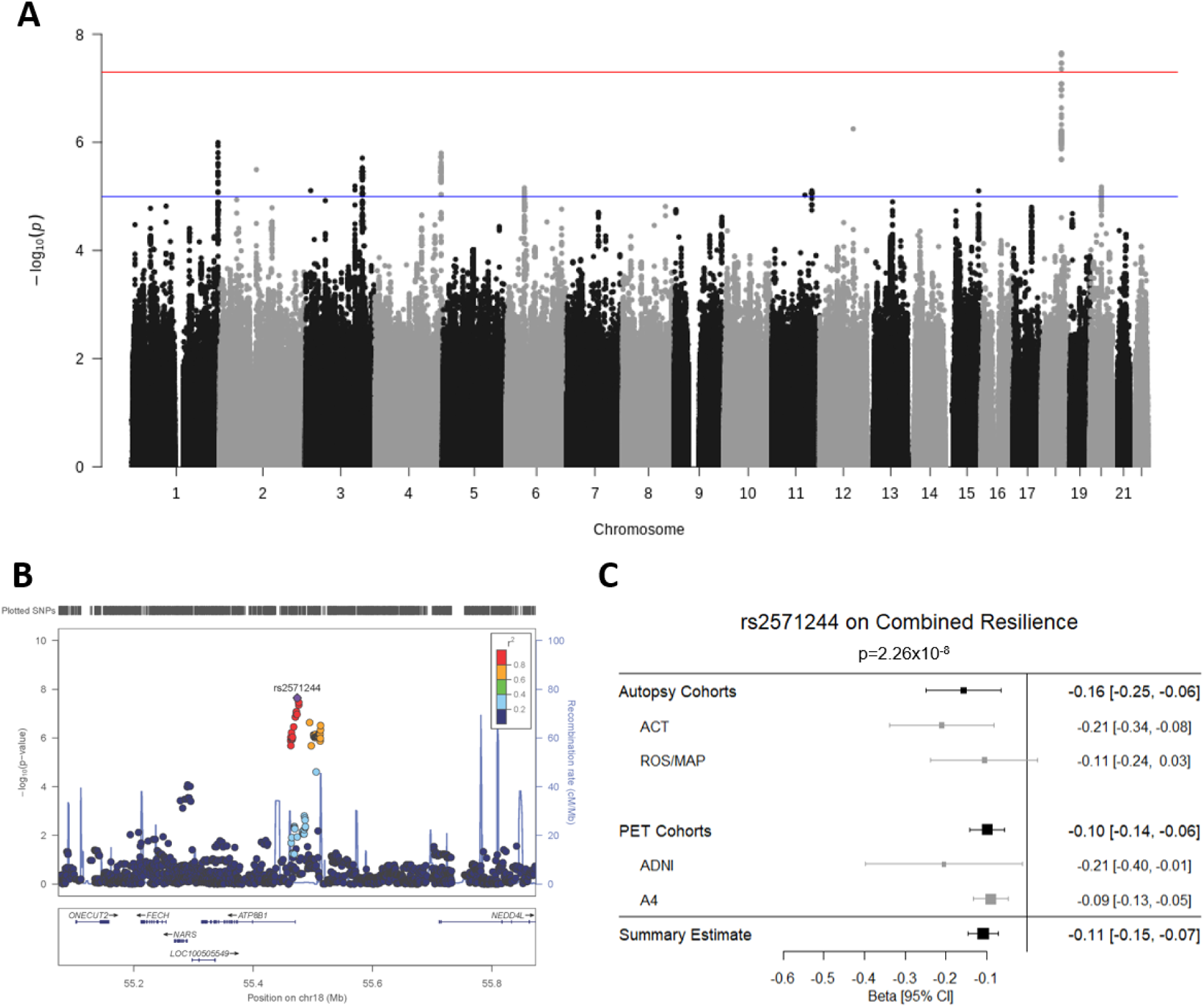
Variant-Level Resilience GWAS Results. **A)** Results from the GWAS analysis of Combined Resilience is presented in a Manhattan Plot. GWAS significance (5×10^−8^) is indicated by the red line, while suggestive significance (1×10^−5^) is indicated by the blue line. **B)** A LocusZoom plot of the GWAS-significant locus on chromosome 18. Colors denote linkage disequilibrium with the most statistically significant SNP. **C)** A forest plot for the top SNP on chromosome 18 is presented demonstrating consistent direction and magnitude of effect across the autopsy and PET datasets and within the component cohorts. The summary estimate at the bottom indicates the meta-analysis of the autopsy and PET combined datasets.

**Figure 5.**
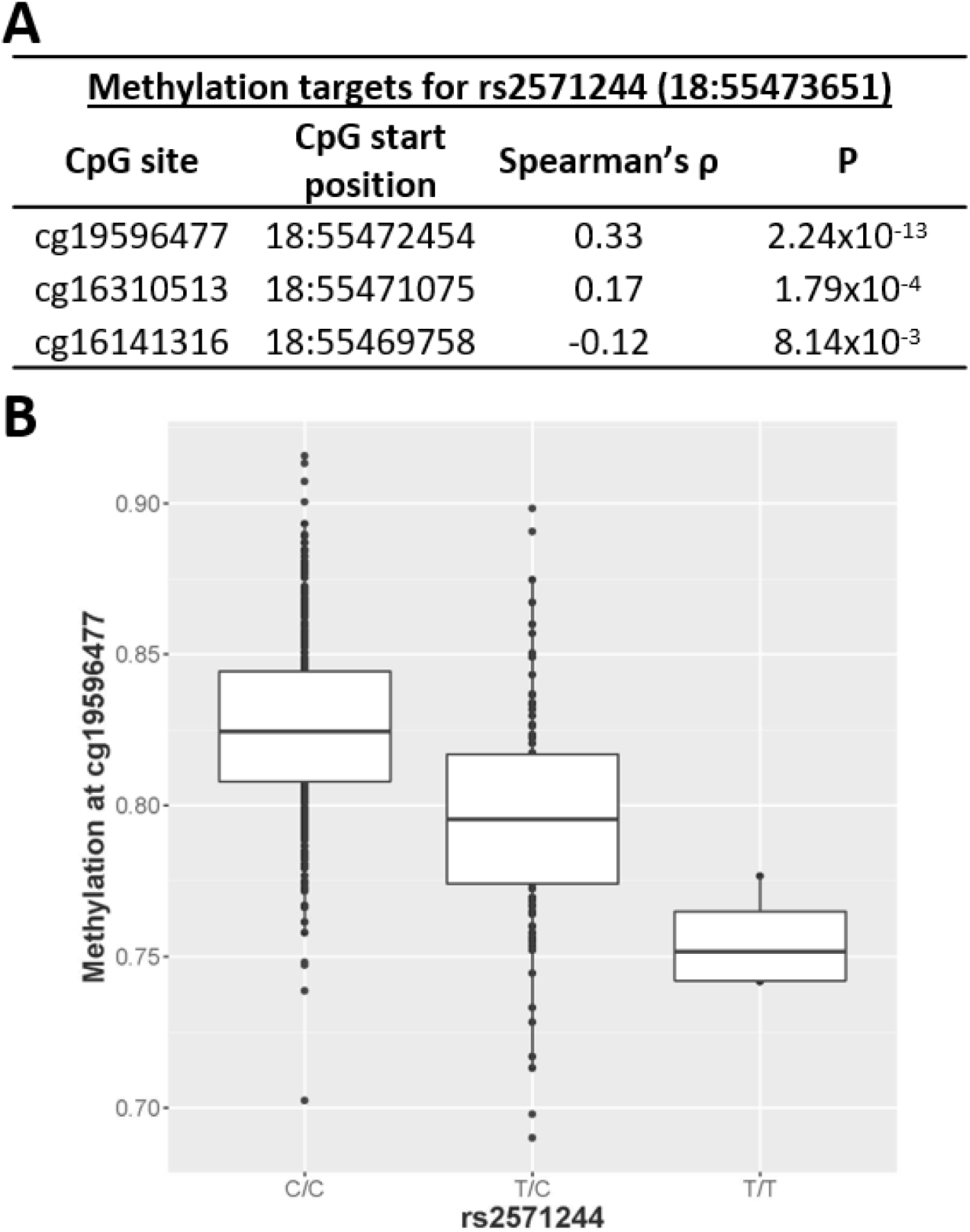
Functional Annotation of Resilience GWAS Results. **A)** The most significant methylation targets for rs2571244 in dorsolateral prefrontal cortex are presented. **B)** The minor allele of rs2571244 (T) is associated with decreased methylation at the CpG site cg19596477.

### Single-Variant Gene Mapping and Functional Annotation

To better characterize the molecular mechanisms of the genome-wide associated loci identified above, we utilized hundreds of functional genomics data sets to test for tissue-specific regulatory activity of these novel variants. The index SNP at the chromosome 18 locus (rs2571244) was strongly associated with prefrontal cortex methylation at multiple sites (**Fig. 4a**) and the minor allele was associated with hypomethylation at a CpG site just upstream of *ATPB81* (cg19596477; p=2×10^−13^; **Fig. 4b**). SNPs in this region also showed statistically significant enrichment for enhancer sites in the Roadmap dataset in across multiple tissues, including brain and liver (adjusted p-values=0.001). However, there was no evidence that rs2571244 functioned as an eQTL or hQTL in any of the databases.

## Discussion

We completed a large genetic analysis of resilience to AD neuropathology and identified a number of variants, genes, and functional pathways that are associated with protection from the downstream consequences of neuropathology. Our results implicate genetic drivers of educational attainment, smoking behaviors, and neuropsychiatric phenotypes in AD resilience; highlight a novel resilience locus on chromosome 18; and implicate metabolism in the liver as a molecular contributor to resilience. Notably, the genetic architecture of resilience appears to be distinct from the genetic architecture of clinical AD, with no observed genetic correlation and nominal contributions of *APOE* on resilience, suggesting that a focus on the molecular contributors to resilience may highlight novel pathways for therapeutic development.

### Resilience Scores are Genetically Correlated with Education, Neuropsychiatric, and Smoking Phenotypes

Results from genetic correlation analyses provided validation of the genetic signals we observed in this analysis and highlighted a number of important biological processes in the etiology of resilience. As expected, we observed strong genetic correlations with educational attainment, cognitive performance, and a number of education-related traits. It is also interesting that we observed some hormone and smoking related traits, although both may be confounded by educational attainment making interpretation challenging. In the case of the smoking traits, genetic risk for smoking and a younger age of initiating smoking was associated with lower levels of resilience, consistent with epidemiological associations between smoking and dementia (Tyas *et al.*, 2003; Peters *et al.*, 2008). In the case of hormone-related phenotypes, an older age of first birth, last birth, and menopause (nominal association in GNOVA and UK Biobank) was correlated with higher resilience scores. Similar associations at the phenotypic level have been reported previously, with an older age at menopause correlated with protection from cognitive decline (Robert N. McLay *et al.*, 2003; Ryan *et al.*, 2009; Ryan *et al.*, 2014). Interestingly, we and others have published extensively on sex differences in the downstream consequences of neuropathology (Buckley *et al.*, 2018; Deming *et al.*, 2018; Hohman *et al.*, 2018; Dumitrescu *et al.*, 2019a; Dumitrescu *et al.*, 2019b). The present results suggest that hormone changes in older adulthood may contribute to susceptibility to cognitive decline, but more work is needed to disentangle the potential contribution of educational attainment on these observed genetic correlations.

In addition, we observed notable genetic correlations with neuropsychiatric phenotypes including ADHD and OCD. Interestingly, genetic risk for OCD was associated with higher resilience scores, while genetic risk for ADHD was associated with lower resilience. Although there is some literature suggesting a potential link between ADHD and dementia, it is challenging because of the symptomatic overlap of the two conditions in adulthood (Callahan *et al.*, 2017). Even less work has characterized the association between OCD and dementia, but the limited literatures suggests OCD is a risk factor for dementia (Dondu *et al.*, 2015). Thus, it is quite interesting that we observe a positive genetic correlation between OCD and resilience here, suggesting a potential protective role. Past work has highlighted a strong negative genetic correlation between OCD and metabolic phenotypes including body mass index, hip circumference, smoking, triglycerides, and insulin levels (Dondu *et al.*, 2015). OCD and ADHD also show a similar opposing genetic correlation with educational attainment, so it may be that the genetic correlation between these psychiatric conditions and resilience is secondary to metabolic or educational attainment phenotypes, but it is an area ripe for future investigation.

### Variants near ATP8B1 are Associated with Resilience

Our top variant level association was observed on Chromosome 18 in relation to the Combined Resilience score that pools information from residual and proxy measures of reserve. The cluster of SNPs associated with Combined Resilience localized just upstream of *ATP8B1*, and the top SNP was robustly associated with methylation at a site also just upstream of *ATP8B1*. Interestingly, prefrontal cortex methylation at this site was strongly associated with Combined Resilience scores in the ROS/MAP dataset, particularly among rs2571244 minor allele carriers, further implicating methylation as a potential biological driver at this locus. *ATP8B1* is a protein coding gene that encodes an aminophospholipid translocase that is critical for maintaining bile acid homeostasis in the liver (Bull *et al.*, 1998). For that reason, we also performed post-hoc analyses using recently quantified metabolomic measures of 15 bile acids from serum samples in ADNI and observed that the variant was nominally associated with five bile acids, including TCA, GLCA, GCA, TDCA, and TCDCA (p<0.05; see **Supplementary Table 16**). Moreover, we observed significant associations between GLCA and TDCA on Combined Resilience, whereby higher levels of these bile acids were associated with lower levels of resilience (**Supplementary Table 17**). Bile acids have emerged as a potential biological contributor to AD, with recent work reporting differential abundance in AD cases compared to controls in both blood and brain (MahmoudianDehkordi *et al.*, 2019), and other work reporting associations with biomarkers of AD neuropathology (Nho *et al.*, 2019). Notably, both GLCA and TDCA were reported to have robust associations with hippocampal atrophy and glucose hypometabolism. The present findings therefore suggest genetic variation that predisposes some individuals towards a more detrimental bile acid state may also increase susceptibility to cognitive decline. The exact causal pathway of such bile acid effects is difficult to infer. Notably, the methylation QTL that we observed for rs2571244 was in prefrontal cortex, suggesting effects could be through brain, but there is a pressing need to better understand the gut-liver-brain axis and determine whether associations with cognitive aging and dementia are driven by metabolic processes in liver, gut, brain, or all three tissues.

### Pathway Analyses Highlight Metabolism

While variant-level results implicate the metabolic processes in the liver, enrichment results highlight the related branched chain amino acid (BCAA) and dehydrogenase molecular pathways. Although the role of BCAAs in AD onset and progression is unclear, several studies have supported a connection. A previous GWAS study showed that SNPs associated with increased isoleucine plasma levels were also associated with AD (Larsson and Markus, 2017). However, metabolomic studies have shown that increased serum concentration of BCAAs are associated with decreased AD risk (Tynkkynen *et al.*, 2018). Particularly, increased serum valine was associated with decreased rates of cerebral atrophy and cognitive decline (Toledo *et al.*, 2017). Deficits in brain BCAA metabolism have been proposed to contribute to the onset and progression of AD in mice, and increased circulating BCAAs have been hypothesized to increase neuronal mTOR signaling, leading to hyperphosphorylated tau pathology (Li *et al.*, 2018).

Several components of dehydrogenase pathways have been implicated to play a role in dysfunctional oxidative stress handling in AD (Martins *et al.*, 1986). Inhibition of alpha-ketoglutarate, pyruvate, and alcohol dehydrogenases by amyloid beta is thought to contribute to mitochondrial and metabolic dysfunction associated with AD (Casley *et al.*, 2002; Yan and Stern, 2005). Alpha-ketoglutarate dehydrogenase complex expression and activity is reduced in the temporal cortex of AD brains and is thought to reduce energy metabolism, contributing to neurodegeneration (Mastrogiacoma *et al.*, 1996).

### Strengths and Limitations

This project has multiple strengths including the large, well characterized cohorts, the deep phenotypic data that allowed for a quantification of residual cognitive performance given level of amyloidosis, and the comprehensive follow-up analyses highlighting novel genes and pathways contributing to resilience. The study is not without limitations. Our sample was restricted to non-Hispanic white individuals who were healthy and highly educated, limiting generalizability beyond such populations.

Additionally, while we were able to fully harmonize cognitive data within the autopsy and PET analyses separately, subtle differences in the scores across autopsy and PET remain possible due to limited availability of item anchors across all cohorts. Further, we were limited to cross-sectional analyses, which leave open the possibility that some individuals will later develop more severe pathology or cognitive impairment. Additional measures of neuropathology, particularly tau and cerebrovascular pathology, may have explained additional variance in cognitive performance and is an important area for future work. Moreover, the lack of extensive neuropsychological protocols in some datasets limited our ability to investigate other cognitive domains (e.g., language or visuospatial abilities). Finally, while this is the largest analysis of the genetic predictors of residual cognition completed to date, we remained underpowered to detect single variant effects, particularly at a low minor allele frequency. Continued efforts to pool, harmonize, and analyze biomarker, autopsy, and neuropsychological data from larger, more representative cohorts will be needed to more fully characterize the genetic architecture of resilience.

## Conclusions

We completed a large analysis of genetic resilience to AD and highlight several novel biological pathways that may protect the brain from the downstream consequences of amyloidosis. Our results implicate genetic drivers of bile acid homeostasis, vascular and metabolic risk factors, and neuropsychiatric conditions in AD resilience.

## Acknowledgements

This research was supported in part by K01-AG049164, R01-AG059716, R21-AG05994, K12-HD043483, K24-AG046373, HHSN311201600276P, S10-OD023680, R01-AG034962, R01-NS100980, R01-AG056534, P30-AG010161, R01-AG057914, R01-AG15819, R01-AG17917, U01-AG061356, U01-AG006781, K99-AG061238, U01-AG46152, Howard Hughes Medical Institute James H. Gilliam Fellowship for Advanced Study (FEC), F31-AG059345 (FEC), UL1-TR000445 and the Vanderbilt Memory & Alzheimer’s Center.

The results published here are in part based on data obtained from the AMP-AD Knowledge Portal (doi:10.7303/syn2580853). MSBB data were generated from postmortem brain tissue collected through the Mount Sinai VA Medical Center Brain Bank and were provided by Dr. Eric Schadt from Mount Sinai School of Medicine. MayoRNAseq data were provided by the following sources: The Mayo Clinic Alzheimer’s Disease Genetic Studies, led by Dr. Nilufer Ertekin-Taner and Dr. Steven G. Younkin, Mayo Clinic, Jacksonville, FL using samples from the Mayo Clinic Study of Aging, the Mayo Clinic Alzheimer’s Disease Research Center, and the Mayo Clinic Brain Bank. Data collection was supported through funding by NIA grants P50-AG016574, P50-AG005136, R01-AG032990, U01-AG046139, R01-AG018023, U01-AG006576, U01-AG006786, R01-AG025711, R01-AG017216, R01 AG003949, NINDS grant R01-NS080820, CurePSP Foundation, and support from Mayo Foundation. Study data includes samples collected through the Sun Health Research Institute Brain and Body Donation Program of Sun City, Arizona. The Brain and Body Donation Program is supported by the National Institute of Neurological Disorders and Stroke (U24-NS072026 National Brain and Tissue Resource for Parkinson’s Disease and Related Disorders), the National Institute on Aging (P30-AG19610 Arizona Alzheimer’s Disease Core Center), the Arizona Department of Health Services (contract 211002, Arizona Alzheimer’s Research Center), the Arizona Biomedical Research Commission (contracts 4001, 0011, 05-901 and 1001 to the Arizona Parkinson’s Disease Consortium) and the Michael J. Fox Foundation for Parkinson’s Research.

Data were generated as part of the CommonMind Consortium supported by funding from Takeda Pharmaceuticals Company Limited, F. Hoffman-La Roche Ltd and NIH grants R01-MH085542, R01-MH093725, P50-MH066392, P50-MH080405, R01-MH097276, RO1-MH075916, P50-M096891, P50-MH084053S1, R37-MH057881, AG02219, AG05138, MH06692, R01-MH110921, R01-MH109677, R01-MH109897, U01-MH103392, and contract HHSN271201300031C through IRP NIMH. Brain tissue for the study was obtained from the following brain bank collections: the Mount Sinai NIH Brain and Tissue Repository, the University of Pennsylvania Alzheimer’s Disease Core Center, the University of Pittsburgh NeuroBioBank and Brain and Tissue Repositories, and the NIMH Human Brain Collection Core. CMC Leadership: Panos Roussos, Joseph Buxbaum, Andrew Chess, Schahram Akbarian, Vahram Haroutunian (Icahn School of Medicine at Mount Sinai), Bernie Devlin, David Lewis (University of Pittsburgh), Raquel Gur, Chang-Gyu Hahn (University of Pennsylvania), Enrico Domenici (University of Trento), Mette A. Peters, Solveig Sieberts (Sage Bionetworks), Thomas Lehner, Stefano Marenco, Barbara K. Lipska (NIMH).

Data collection and sharing for this project was funded by the Alzheimer’s Disease Neuroimaging Initiative (ADNI) (National Institutes of Health Grant U01 AG024904) and DOD ADNI (Department of Defense award number W81XWH-12-2-0012). ADNI is funded by the National Institute on Aging, the National Institute of Biomedical Imaging and Bioengineering, and through generous contributions from the following: AbbVie, Alzheimer’s Association; Alzheimer’s Drug Discovery Foundation; Araclon Biotech; BioClinica, Inc.; Biogen Inc. Cambridge, MA 02139, provided support for genotyping of the A4 Study cohort; Bristol-Myers Squibb Company; CereSpir, Inc.; Cogstate; Eisai Inc.; Elan Pharmaceuticals, Inc.; Eli Lilly and Company; EuroImmun; F. Hoffmann-La Roche Ltd and its affiliated company Genentech, Inc.; Fujirebio; GE Healthcare; IXICO Ltd.;Janssen Alzheimer Immunotherapy Research & Development, LLC.; Johnson & Johnson Pharmaceutical Research & Development LLC.; Lumosity; Lundbeck; Merck & Co., Inc.;Meso Scale Diagnostics, LLC.; NeuroRx Research; Neurotrack Technologies; Novartis Pharmaceuticals Corporation; Pfizer Inc.; Piramal Imaging; Servier; Takeda Pharmaceutical Company; and Transition Therapeutics. The Canadian Institutes of Health Research is providing funds to support ADNI clinical sites in Canada. Private sector contributions are facilitated by the Foundation for the National Institutes of Health (www.fnih.org). The grantee organization is the Northern California Institute for Research and Education, and the study is coordinated by the Alzheimer’s Therapeutic Research Institute at the University of Southern California. ADNI data are disseminated by the Laboratory for Neuro Imaging at the University of Southern California.

Additional data collection and sharing for this project was funded by the Alzheimer’s Disease Metabolomics Consortium (National Institute on Aging R01-AG046171, RF1-AG051550 and 3U01-AG024904-09S4).

## Conflicts of Interest

Nothing to report.

## References

Amlie-Wolf A, Tang M, Mlynarski EE, Kuksa PP, Valladares O, Katanic Z, et al. INFERNO: inferring the molecular mechanisms of noncoding genetic variants. Nucleic Acids Research 2018; 46(17): 8740–53.

Bennett DA, Buchman AS, Boyle PA, Barnes LL, Wilson RS, Schneider JA. Religious Orders Study and Rush Memory and Aging Project. Journal of Alzheimer’s Disease 2018(Preprint): 1–28.

Boyle PA, Yu L, Leurgans SE, Wilson RS, Brookmeyer R, Schneider JA, et al. Attributable risk of Alzheimer’s dementia attributed to age-related neuropathologies. Ann Neurol 2019; 85(1): 114–24.

Buckley RF, Mormino EC, Amariglio RE, Properzi MJ, Rabin JS, Lim YY, et al. Sex, amyloid, and APOE ε4 and risk of cognitive decline in preclinical Alzheimer’s disease: Findings from three well-characterized cohorts. Alzheimer’s & Dementia 2018.

Bull LN, van Eijk MJ, Pawlikowska L, DeYoung JA, Juijn JA, Liao M, et al. A gene encoding a P-type ATPase mutated in two forms of hereditary cholestasis. Nat Genet 1998; 18(3): 219–24.

Callahan BL, Bierstone D, Stuss DT, Black SE. Adult ADHD: Risk Factor for Dementia or Phenotypic Mimic? Front Aging Neurosci 2017; 9: 260.

Casley CS, Canevari L, Land JM, Clark JB, Sharpe MA. Beta-amyloid inhibits integrated mitochondrial respiration and key enzyme activities. J Neurochem 2002; 80(1): 91–100.

Crane PK, Trittschuh E, Mukherjee S, Saykin AJ, Sanders RE, Larson EB, et al. Incidence of cognitively defined late-onset Alzheimer’s dementia subgroups from a prospective cohort study. Alzheimers Dement 2017; 13(12): 1307–16.

Deming Y, Dumitrescu L, Barnes LL, Thambisetty M, Kunkle B, Gifford KA, et al. Sex-specific genetic predictors of Alzheimer’s disease biomarkers. Acta neuropathologica 2018: 1–16.

Dondu A, Sevincoka L, Akyol A, Tataroglu C. Is obsessive-compulsive symptomatology a risk factor for Alzheimer-type dementia? Psychiatry Res 2015; 225(3): 381–6.

Dumitrescu L, Barnes LL, Thambisetty M, Beecham G, Kunkle B, Bush WS, et al. Sex differences in the genetic predictors of Alzheimer’s pathology. Brain 2019a; 142(9): 2581–9.

Dumitrescu L, Mayeda ER, Sharman K, Moore AM, Hohman TJ. Sex difference in the genetic architecture of Alzheimer’s disease. Current Genetic Medicine Reports 2019b.

Franzmeier N, Ren J, Damm A, Monté-Rubio G, Boada M, Ruiz A, et al. The BDNF Val66Met SNP modulates the association between beta-amyloid and hippocampal disconnection in Alzheimer’s disease. Molecular Psychiatry 2019: 1.

Gamazon ER, Zwinderman AH, Cox NJ, Denys D, Derks EM. Multi-tissue transcriptome analyses identify genetic mechanisms underlying neuropsychiatric traits. Nat Genet 2019; 51(6): 933–40.

Hohman TJ, Dumitrescu L, Barnes LL, Thambisetty M, Beecham GW, Kunkle B, et al. Sex-specific effects of Apolipoprotein E with cerebrospinal fluid levels of tau. JAMA Neurology 2018.

Hohman TJ, Dumitrescu L, Cox NJ, Jefferson AL. Genetic resilience to amyloid related cognitive decline. Brain Imaging and Behavior 2016a: 1–9.

Hohman TJ, Koran MEI, Thornton-Wells TA, Initiative AsN. Genetic variation modifies risk for neurodegeneration based on biomarker status. Frontiers in Aging Neuroscience 2014a; 6.

Hohman TJ, Koran MI, Thornton-Wells TA. Genetic Modification of the Relationship between Phosphorylated Tau and Neurodegeneration. Alzheimer’s & Dementia 2014b; 10(6): 637–45.

Hohman TJ, McLaren DG, Mormino EC, Gifford KA, Libon DJ, Jefferson AL. Asymptomatic Alzheimer disease: Defining resilience. Neurology 2016b; 87(23): 2443–50.

Jansen IE, Savage JE, Watanabe K, Bryois J, Williams DM, Steinberg S, et al. Genome-wide meta-analysis identifies new loci and functional pathways influencing Alzheimer’s disease risk. Nature Genetics 2019; 51(3): 404–13.

Kukull WA, Higdon R, Bowen JD, McCormick WC, Teri L, Schellenberg GD, et al. Dementia and Alzheimer disease incidence: a prospective cohort study. Archives of neurology 2002; 59(11): 1737–46.

Kunkle BW, Grenier-Boley B, Sims R, Bis JC, Damotte V, Naj AC, et al. Genetic meta-analysis of diagnosed Alzheimer’s disease identifies new risk loci and implicates Aβ, tau, immunity and lipid processing. Nature Genetics 2019; 51(3): 414–30.

Lambert JC, Ibrahim-Verbaas CA, Harold D, Naj AC, Sims R, Bellenguez C, et al. Meta-analysis of 74,046 individuals identifies 11 new susceptibility loci for Alzheimer’s disease. Nat Genet 2013; 45(12): 1452–8.

Larsson SC, Markus HS. Branched-chain amino acids and Alzheimer’s disease: a Mendelian randomization analysis. Sci Rep 2017; 7(1): 13604.

Li H, Ye D, Xie W, Hua F, Yang Y, Wu J, et al. Defect of branched-chain amino acid metabolism promotes the development of Alzheimer’s disease by targeting the mTOR signaling. Biosci Rep 2018; 38(4).

Liu JZ, Mcrae AF, Nyholt DR, Medland SE, Wray NR, Brown KM, et al. A versatile gene-based test for genome-wide association studies. The American Journal of Human Genetics 2010; 87(1): 139–45.

Lu Q, Li B, Ou D, Erlendsdottir M, Powles RL, Jiang T, et al. A Powerful Approach to Estimating Annotation-Stratified Genetic Covariance via GWAS Summary Statistics. The American Journal of Human Genetics 2017; 101(6): 939–64.

Mägi R, Morris AP. GWAMA: software for genome-wide association meta-analysis. BMC bioinformatics 2010; 11(1): 288.

MahmoudianDehkordi S, Arnold M, Nho K, Ahmad S, Jia W, Xie G, et al. Altered bile acid profile associates with cognitive impairment in Alzheimer’s disease-An emerging role for gut microbiome. Alzheimers Dement 2019; 15(1): 76–92.

Martins RN, Harper CG, Stokes GB, Masters CL. Increased cerebral glucose-6-phosphate dehydrogenase activity in Alzheimer’s disease may reflect oxidative stress. J Neurochem 1986; 46(4): 1042–5.

Mastrogiacoma F, Lindsay JG, Bettendorff L, Rice J, Kish SJ. Brain protein and alpha-ketoglutarate dehydrogenase complex activity in Alzheimer’s disease. Ann Neurol 1996; 39(5): 592–8.

Mirra SS, Heyman A, McKeel D, Sumi SM, Crain BJ, Brownlee LM, et al. The Consortium to Establish a Registry for Alzheimer’s Disease (CERAD) Part II. Standardization of the neuropathologic assessment of Alzheimer’s disease. Neurology 1991; 41(4): 479-.

Mishra A, Macgregor S. VEGAS2: software for more flexible gene-based testing. Twin Research and Human Genetics 2015; 18(01): 86–91.

Mishra A, MacGregor S. A Novel Approach for Pathway Analysis of GWAS Data Highlights Role of BMP Signaling and Muscle Cell Differentiation in Colorectal Cancer Susceptibility. Twin Res Hum Genet 2017; 20(1): 1–9.

Monsell SE, Mock C, Fardo DW, Bertelsen S, Cairns NJ, Roe CM, et al. Genetic Comparison of Symptomatic and Asymptomatic Persons With Alzheimer Disease Neuropathology. Alzheimer Disease & Associated Disorders 2017; 31(3): 232–8.

Monsell SE, Mock C, Roe CM, Ghoshal N, Morris JC, Cairns NJ, et al. Comparison of symptomatic and asymptomatic persons with Alzheimer disease neuropathology. Neurology 2013; 80(23): 2121–9.

Mostafavi S, Gaiteri C, Sullivan SE, White CC, Tasaki S, Xu J, et al. A molecular network of the aging human brain provides insights into the pathology and cognitive decline of Alzheimer’s disease. Nat Neurosci 2018; 21(6): 811–9.

Muthén LK, Muthén BO. Mplus User’s Guide. Seventh ed. Los Angeles, CA: Muthén & Muthén; 1998-2015.

Ng B, White CC, Klein H-U, Sieberts SK, McCabe C, Patrick E, et al. An xQTL map integrates the genetic architecture of the human brain’s transcriptome and epigenome. Nature neuroscience 2017; 20(10): 1418.

Nho K, Kueider-Paisley A, MahmoudianDehkordi S, Arnold M, Risacher SL, Louie G, et al. Altered bile acid profile in mild cognitive impairment and Alzheimer’s disease: Relationship to neuroimaging and CSF biomarkers. Alzheimers Dement 2019; 15(2): 232–44.

Peters R, Poulter R, Warner J, Beckett N, Burch L, Bulpitt C. Smoking, dementia and cognitive decline in the elderly, a systematic review. BMC geriatrics 2008; 8: 36.

Properzi MJ, Buckley RF, Chhatwal JP, Donohue MC, Lois C, Mormino EC, et al. Nonlinear Distributional Mapping (NoDiM) for harmonization across amyloid-PET radiotracers. Neuroimage 2019; 186: 446–54.

Raghavan NS, Dumitrescu L, Mormino E, Mahoney E, Lee AJ, Gao Y, et al. Common Variants in RBFOX1 are Associated with Brain Amyloidosis. In Review 2020.

Rahimi J, Kovacs GG. Prevalence of mixed pathologies in the aging brain. Alzheimers Research & Therapy 2014; 6(9): 82.

Reed BR, Mungas D, Farias ST, Harvey D, Beckett L, Widaman K, et al. Measuring cognitive reserve based on the decomposition of episodic memory variance. Brain 2010: awq154.

Robert N. McLay, Ph.D., M.D.,, Pauline M. Maki, Ph.D., and, Constantine G. Lyketsos, M.D., M.H.S. Nulliparity and Late Menopause Are Associated With Decreased Cognitive Decline. The Journal of Neuropsychiatry and Clinical Neurosciences 2003; 15(2): 161–7.

Ryan J, Carriere I, Scali J, Ritchie K, Ancelin ML. Life-time estrogen exposure and cognitive functioning in later life. Psychoneuroendocrinology 2009; 34(2): 287–98.

Ryan J, Scali J, Carriere I, Amieva H, Rouaud O, Berr C, et al. Impact of a premature menopause on cognitive function in later life. Bjog 2014; 121(13): 1729–39.

Sonnen JA, Santa Cruz K, Hemmy LS, Woltjer R, Leverenz JB, Montine KS, et al. Ecology of the aging human brain. Archives of Neurology 2011; 68(8): 1049–56.

Sperling RA, Rentz DM, Johnson KA, Karlawish J, Donohue M, Salmon DP, et al. The A4 study: stopping AD before symptoms begin? Science translational medicine 2014; 6(228): 228fs13–fs13.

Toledo JB, Arnold M, Kastenmuller G, Chang R, Baillie RA, Han X, et al. Metabolic network failures in Alzheimer’s disease: A biochemical road map. Alzheimers Dement 2017; 13(9): 965–84.

Tyas SL, White LR, Petrovitch H, Webster Ross G, Foley DJ, Heimovitz HK, et al. Mid-life smoking and late-life dementia: the Honolulu-Asia Aging Study. Neurobiol Aging 2003; 24(4): 589–96.

Tynkkynen J, Chouraki V, van der Lee SJ, Hernesniemi J, Yang Q, Li S, et al. Association of branched-chain amino acids and other circulating metabolites with risk of incident dementia and Alzheimer’s disease: A prospective study in eight cohorts. Alzheimers Dement 2018; 14(6): 723–33.

White CC, Yang H-S, Yu L, Chibnik LB, Dawe RJ, Yang J, et al. Identification of genes associated with dissociation of cognitive performance and neuropathological burden: Multistep analysis of genetic, epigenetic, and transcriptional data. PLoS medicine 2017; 14(4): e1002287.

Yan D, Hu B, Darst BF, Mukherjee S, Kunkle BW, Deming Y, et al. Biobank-wide association scan identifies risk factors for late-onset Alzheimer’s disease and endophenotypes. 2018: 468306.

Yan SD, Stern DM. Mitochondrial dysfunction and Alzheimer’s disease: role of amyloid-beta peptide alcohol dehydrogenase (ABAD). Int J Exp Pathol 2005; 86(3): 161–71.

Yang J, Lee SH, Goddard ME, Visscher PM. GCTA: a tool for genome-wide complex trait analysis. Am J Hum Genet 2011; 88(1): 76–82.

Yu L, Boyle PA, Segawa E, Leurgans S, Schneider JA, Wilson RS, et al. Residual decline in cognition after adjustment for common neuropathologic conditions. Neuropsychology 2015; 29(3): 335–43.

Yu L, Petyuk VA, Gaiteri C, Mostafavi S, Young-Pearse T, Shah RC, et al. Targeted brain proteomics uncover multiple pathways to Alzheimer’s dementia. Ann Neurol 2018; 84(1): 78–88. Genetic Markers of Resilience 33

