## Supplemental Methods for "Genetic Variants and Functional Pathways Associated with Resilience to Alzheimer’s Disease"

### Supplemental Text 1. Psychometric Analyses.

#### 1A. Confirmatory Factor Analyses in Each Study.

**Step 1: Domain assignment:** In each of the studies (Adult Changes in Thought [ACT], Alzheimer’s Disease Neuroimaging Initiative [ADNI], the Religious Orders Study–Memory and Aging Project (ROS/MAP], and the Anti-Amyloid Treatment in Asymptomatic Alzheimer's study [A4]), the expert panel (Dr. Trittschuh, Dr. Mez, and Dr. Saykin) assigned items from the neuropsychological battery to one of the four domains (memory, language, executive functioning, and visuospatial ability); other items did not map to any of these domains. For the present analyses, we completed harmonization for memory and executive function.

**Step 2: Data quality control:** Quality control was performed for each study. To maximize representation across diagnostic categories, we prepared a data set which included item-level data for individuals at their final visit. Before running psychometric models, we performed additional recoding of the data to make sure lower values represented lower cognitive performance (e.g., Trails A and B were reverse coded). We also considered the distribution of each item among those with non-missing data and combined categories as needed. Our goals were a.) to avoid sparse categories (operationally defined as <5 responses for each study administering each item) and b.) to have a maximum of 10 categories, which is the maximum number of categories handled by Mplus v7.4 [2]. We treated each item as an ordinal indicator of the domain—the numerical value assigned to each category is irrelevant beyond its rank.

We also looked at informative missingness in each study and recoded relevant items accordingly. For example, some of the studies include multiple missing codes, where it was possible to identify refusal to respond to an item as opposed to the interviewer ran out of time and the item was never administered. The first of these—refusal—we took as informative missing and assigned that code to the lowest response category, while the second of these—missing due to scheduling etc.—we took as non-informative missing and omitted that item from consideration.

**Step 3: Confirmatory factor analyses:** We used cognitive data from individual’s last visit for running psychometric analyses to derive a robust composite which captures the most variance in a given neuropsychological battery. We then turned to confirmatory factor analysis (CFA) modeling with Mplus using Robust Weighted Least Squares including terms for the mean and the variance (WLSMV) estimator. We ran two models: a.) a single factor model, with no residual structure; and b.) a data-driven bifactor model, using hierarchical clustering-assigned sub-domains. We consulted the expert panel on the sub-domain assignment of items in our data-driven approach to make sure these models made sense to our experts. Our overall strategy was that we would choose the single factor model if adding secondary factors did not markedly improve model fit and if adding secondary factors did not markedly impact any individual’s score (see below).

Our criteria for selecting the best model included fit statistics (see below) and concordance of model results with theory, such as all loadings on secondary factors being positive. The fit statistics we considered were the confirmatory fit index (CFI), where higher values indicate better fit, and thresholds of 0.90 and 0.95 have been used in other settings as criteria for adequate or good fit [1, 3]; the Tucker-Lewis Index (TLI), which has similar criteria as the CFI; and the root mean squared error of approximation (RMSEA), where lower values indicate better fit, and thresholds of 0.08 and 0.05 have been used in other settings as criteria for adequate or good fit [1, 3].

When comparing the single factor model with the best bifactor model, we a.) looked at whether loadings on the primary factor were within 10% of each other across the two models and b.) compared the scores for the single factor model vs. scores for the candidate bifactor model. Our threshold was a difference of 0.30 units, which was based on the default stopping rule for computerized adaptive testing; this has been used for years as the default level of tolerable measurement differences in the setting of computerized adaptive tests. While arbitrary, this is a level of ambiguity that has been thought to be tolerable in a variety of situations. If there were a substantial number of people for whom the differences in scores were larger than 0.3 from each other, and if the bifactor model had better fit statistics, we selected the bifactor model for modeling a domain.

#### 1B. Co-calibration of the Domains Across ACT, ADNI, and ROS/MAP

**Step 1: Identification of anchor items:** Co-calibration requires either the same people taking different tests or different tests sharing common items. Here we had common items. We identified candidate anchor items with identical content across tests administered in different studies and ensured that their relationship with the underlying ability tested was the same across studies by performing preliminary confirmatory factor analysis models within each study. These items were then used to anchor the scales in each domain to a common metric**.** We consulted a member of the expert panel (Dr. Trittschuh) to make sure we chose the anchor items correctly.

**Step 2: Quality control for anchor items:** Anchor items were cleaned and recoded after merging items from all the studies, making sure that the ranges of the anchor items were similar in each study. We carefully reviewed documentation from each study to ensure that the stimulus was precisely the same, that the response options were precisely the same or could be re-coded to be the same, and that we were mapping data from each study in a way that the same response would result in the same score regardless of which study the person was enrolled in.

A note regarding response options—in many cases the stimulus is fairly open-ended, such as “can you please draw from memory the figure you copied a while ago”, where the participant is handed a blank sheet of paper and a writing implement. The resulting drawing then gets scored based on how similar it was to the initial stimulus figure. The specific scoring applied to such a stimulus could vary across studies. One study could score such an item as correct vs. incorrect, while another could apply points for various aspects of the drawing. In this instance, we would review the first study’s scoring documentation to determine how many aspects of the drawing needed to be present for a “correct” score. Then we would map all scores from the second study that would have resulted in a “correct” or “incorrect” score in the first study. In this way, the resulting score is invariant to which study the person is participating in, as each response would be consistently scored regardless of study.

**Step 3: Confirmatory factor analyses:** We co-calibrated memory and executive function by incorporating the components of the best model in each study (i.e., the final single-factor or bifactor model selected as described above) into one mega-calibration model.

One particularly tricky aspect of co-calibrating scores using bifactor models is how to handle secondary domains. Some anchor items had loadings on the primary domain (e.g. memory) and also on a secondary domain. That structure by itself does not lead to conceptual problems. However, item representation of the secondary domain may vary across studies, with variable numbers of items, and potential missing data and identifiability issues. To address this, we used robust maximum likelihood (MLR) estimation that is robust to missing data and assigned all subdomain indicators across studies to the same subdomain. Unlike running a CFA model with the WLSMV estimator, a CFA model with MLR estimator does not output fit statistics like CFI/TLI/RMSEA. For our purposes, these secondary domains were nuisances. We performed a number of sensitivity analyses to reassure ourselves that scores on the primary domain were minimally impacted by various ways of specifying the mean and variance on secondary domains. In the final models, we specified a mean of 0 and a variance of 1 for each secondary domain factor, regardless of the number of studies that included items that loaded on that factor.

Once we had fit the final mega-calibration model for each domain, we extracted factor scores for the primary factor (e.g. memory). The resulting scores are on the same metric with a mean of 0 and variance of 1. We used all participants with relevant data to fit data for each domain, so the scale for each domain was based on models that included different people, since some people were missing for some domains. Therefore, we used ACT as a reference population for standardizing scores for each domain, as it is a community-based prospective cohort study with a very large sample (n=825) of people with sufficient cognitive data. We applied the same standardization to all participants for each study. Thus, a score of 0, regardless of study, reflects the mean for people in their last visit in the ACT study; and a score of -1, regardless of study, reflects 1 standard deviation below the mean for people in the ACT study.

For A4, we used these estimated thresholds and loadings of items from the co-calibration mega-calibration models to obtain scores for individuals. New items (not part of ACT, ADNI, and ROS/MAP) were freely estimated while already seen items will have their parameters fixed based on these mega-calibration models.

#### 1C. Confirmatory Factor Analyses Model Considerations in Co-calibration Models

**1.** For all CFA models, we categorized items to ≤ 10 categories. For co-calibration purpose, we had to re-categorize some of the items even though they already had ≤ 10 categories. This was because some studies had more granular data (more categories) for anchor items compared to other studies. In these cases, after we estimated item parameters from the co-calibration model, we re-estimated parameters of the anchor item(s) in the most granular form in the given study.

After using re-coded items for co-calibration, we fixed all of the other items to their values from the co-calibration run and freely estimated parameters for re-coded anchors in their most granular form. This approach enabled us to obtain more precise scores in studies that incorporated more granular scoring rules, while still using all items administered across studies to co-calibrate metrics across studies.

**2.** The base co-calibration exercise for each of the four domains was performed across ACT, ROS/MAP, and ADNI. The A4 data were subsequently added with the following steps. For memory, we identified anchor items and fixed their item parameters to those estimated previously in the base co-calibration models; unique items administered in the new study that were not administered to people in ACT, ROS/MAP, or ADNI were freely estimated.

In these models:

a) The mean and variance for the primary factor were freely estimated.

b) If every item in a sub-domain in the new data had parameters available from the co-calibration model, we fixed those item parameters to their previously identified values and allowed the mean and variance to be freely estimated in the new data.

c) If no item from a sub-domain had parameters available, then we freely estimated each of the sub-domain loadings, fixing the mean and variance of the subdomain factor to 0 and 1.

d) If there was a mix of previously specified and new items in a subdomain, we fixed the parameters for the previously specified items and allowed the mean and variance of the factor and the loadings for new items to be freely estimated in the new data.

**NOTE:** A more detailed overview and all code snippets can be obtained from authors on request.

### Supplemental Text 2. Residual Models of Resilience

#### 2A. Quantifying Residuals for Inclusion in Latent Variable Models of Resilience

**Cognitive Measures.** The harmonized metrics for memory and executive function outlined above were included as outcome measures. As seen in **Figure S1,** the harmonized metrics for memory performed quite well across datasets and cognitively normal participants showed comparable distributions across cohorts. However, the A4 dataset did not contain sufficient anchor items to harmonize executive function. As an additional cognitive item, we leveraged the preclinical cognitive composite measure that had been previously quantified in A4, and quantified an equivalent metric in ADNI. As can be seen in **Figure S2**, the 4-item PACC was well anchored across datasets and behaved comparably in the normal cognition participants across both studies. Similarly, the calibrated memory overlaid quite well across all four datasets.

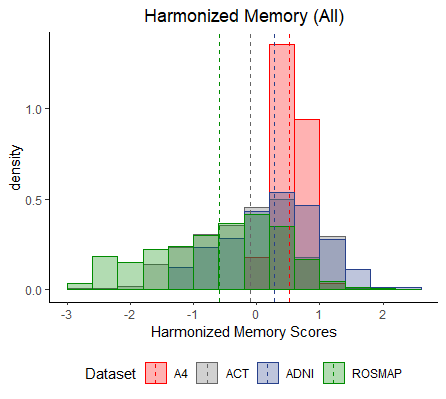

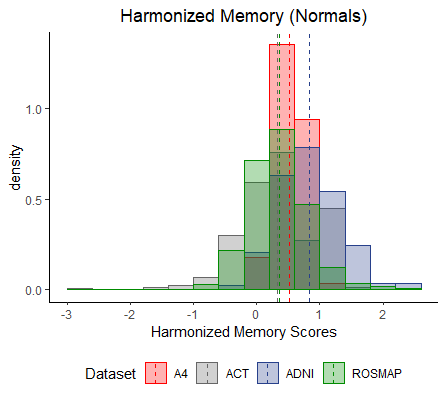

**Figure S1. Harmonized Memory Measures by Dataset.** The harmonized memory measure is presented on the x-axis, and the density of observations at a given value is presented on the y-axis. Histograms are colored by dataset. All participants are presented in the left panel, and cognitively normal participants are presented in the right panel.

**Calculating Residuals in PET Datasets.** Our primary analytical model included individuals across the diagnostic spectrum from normal cognition to Alzheimer’s disease. The A4 dataset only includes individuals with normal cognition, and for that reason we used harmonized measures of amyloid and cognition to build our models across the ADNI and A4 dataset so that linear regression models reflected the strong association between amyloid levels and cognition among participants with mild cognitive impairment (MCI) and AD. Each of the cognitive scores was regressed on age, sex, and our harmonized measure of amyloid. We ran the models in the whole sample as well as in a subset of cognitively normal individuals. See the **Table** for the variance explained by each of the models. Residuals from these individual linear regression models were stored to act as indicators in Latent Variable Models specified in **2B** below.

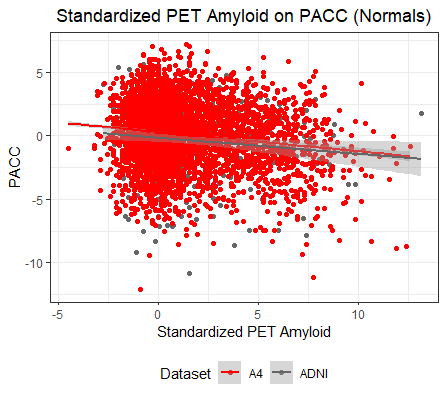

**Figure S2. Amyloid association with cognition in ADNI and A4.** The standardized PET amyloid metric is presented on the x-axis, the Preclinical Alzheimer’s Cognitive Composite (PACC) is presented on the y-axis. The points and lines are colored by dataset.

**Calculating Residuals in Autopsy Datasets.** Similar to our approach in PET datasets, we pooled data across the ROS/MAP and ACT cohort leveraging harmonized measures of amyloid and cognitive performance. As above, cognition was regressed on age at death, sex, postmortem interval, and CERAD staging (treated as an ordinal variable). Variance explained by each of the models in the whole sample and in the subset of cognitively normal individuals is presented in the **Table**. Residuals from individual linear regression models were stored to input as indicator variables in latent variable models specified in **2B** below.

**Table. Variance Explained**

| **Outcome** | **Dataset** | **Variance Explained (R^2^)**  **Whole Sample** | **Variance Explained (R^2^)**  **Normal Cognition** |
| --- | --- | --- | --- |
| Executive Function | Autopsy | 13% | 6% |
|  | PET (ADNI only) | 17% | 13% |
| Memory | Autopsy | 20% | 7% |
|  | PET | 19% | 20% |
| PACC | PET | 20% | 15% |

#### 2B. Calculating a Latent Variable Model of Resilience

A confirmatory factor analysis was performed in Mplus (version 7.3) using the WLSMV estimator. Residuals from models that included overlapping cognitive items were treated as anchors across datasets to ensure the final resilience phenotypes were on a comparable scale across datasets. The model diagram is presented in **Figure 1** in the main text of the manuscript. All indicator variables were treated as continuous variables in all models. The factor scores were extracted from the model and leveraged as outcome variables in our GWAS analyses. Separate models were calculated using residuals from the entire sample and residuals from the cognitive normal subset of participants. Model fit for the overall sample was good (χ^2^=107, RMSEA=0.07, CFI=0.98, TLI=0.96). The model fit in the cognitively normal subset was similar and good (χ^2^=48, RMSEA=0.05, CFI=0.99, TLI=0.98). Although the model fits were similar, the final resilience metrics behaved much differently in the A4 dataset, depending on whether or not individuals with MCI and AD were included in the original regression analyses. It is unclear whether the most appropriate model should include the association between amyloid levels and cognitive performance among participants with MCI and AD, or whether it is more appropriate to restrict the sample to cognitively normal participants, so we ran analyses both ways and present results for both approaches in the main body of the manuscript. Needless to say, restricting to cognitively normal participants reduced heterogeneity across studies and, thus, may have improved statistical power in meta-analyses despite the reduction in sample size.

### Supplemental Text 3. Detailed Tables on the Neuropsychological Items by Domain for Each Study and Fit Statistics from Harmonization Analyses

#### 3A. Memory

**ACT:** Final model was a data driven methods-effects bifactor model with CFI = 0.951, TLI = 0.946, and RMSEA = 0.065. The following items were included in the CFA analysis:

| **Study** | **Variable** | **Description** | **Secondary Structure** |
| --- | --- | --- | --- |
| ACT | mat_mem | Mattis Dementia Rating Scale Memory score | F1 |
| ACT | w_in_c1 | Word list learning trial 1 total score | F1 |
| ACT | w_in_c2 | Word list learning trial 2 total score | F1 |
| ACT | w_in_c3 | Word list learning trial 3 total score | F1 |
| ACT | w_rcl_c | Word List Recall—correct | F1 |
| ACT | w_rcg_t | Word Recognition—total correct | F1 |
| ACT | cp_re_ci | Constructional Praxis Delay—circle |  |
| ACT | cp_re_di | Constructional Praxis Delay—diamond |  |
| ACT | cp_re_re | Constructional Praxis Delay—rectangles |  |
| ACT | cp_re_cu | Constructional Praxis Delay—cube |  |
| ACT | w_lm_ima | Logical Mem I—immediate recall total story A | F2 |
| ACT | w_lm_imb | Logical Mem I—immediate recall total story B | F3 |
| ACT | w_lm_dea | Logical Mem II—delayed recall total story A | F2 |
| ACT | w_lm_deb | Logical Mem II—delayed recall total story B | F3 |
| ACT | w_vp_ine | Verbal Paired Associates I easy | F4 |
| ACT | w_vp_inh | Verbal Paired Associates I hard | F5 |
| ACT | w_vp_ree | Verbal Paired Associates II easy | F4 |
| ACT | w_vp_reh | Verbal Paired Associates II hard | F5 |
| ACT-CASI | rgs1 | repeat words |  |
| ACT-CASI | rc1a | Word recall—something to wear—1 | F6 |
| ACT-CASI | rc1b | Word recall—a color—1 | F6 |
| ACT-CASI | rc1c | Word recall—personal quality—1 | F6 |
| ACT-CASI | yr | What is today’s date?—year |  |
| ACT-CASI | mo | What is today’s date?—month | F7 |
| ACT-CASI | casi_dat | What is today’s date?—day | F7 |
| ACT-CASI | day | What day of week? |  |
| ACT-CASI | casi_ssn | What season is it? |  |
| ACT-CASI | spa | What state and city? |  |
| ACT-CASI | spb | What is this place? |  |
| ACT-CASI | rc2a | Word recall—something to wear—2 | F6 |
| ACT-CASI | rc2b | Word recall—a color—2 | F6 |
| ACT-CASI | rc2c | Word recall—personal quality—2 | F6 |
| ACT-CASI | rcobj | Recall of 5 objects |  |

**ADNI:** Final model was a data driven bifactor model with CFI = 0.984, TLI = 0.982, and RMSEA = 0.084. The following items were included in the CFA analysis:

| **Study** | **Variable** | **Description** | **Secondary Structure** |
| --- | --- | --- | --- |
| ADNI | limmtotal | Logical Memory—Immediate Recall | F1 |
| ADNI | ldeltotal | Logical Memory—Delayed Recall | F1 |
| ADNI | avtot1* | AVLT Trial 1 Total |  |
| ADNI | avtot2* | Trial 2 Total | F2 |
| ADNI | avtot3* | Trial 3 Total | F2 |
| ADNI | avtot4* | Trial 4 Total | F2 |
| ADNI | avtot5* | Trial 5 Total | F2 |
| ADNI | avtot6* | Trial 6 Total | F2 |
| ADNI | avtotb* | List B Total |  |
| ADNI | avdel30min* | 30 Minute Delay Total | F2 |
| ADNI | avdeltot* | Recognition Score |  |
| ADNI | q1score | ADAS Word Recall—score | F3 |
| ADNI | q4score | ADAS Delayed Word Recall | F3 |
| ADNI | q7score | ADAS Orientation—score |  |
| ADNI | q8score | ADAS Word Recognition—score |  |
| ADNI | mmdate | What is today's date? |  |
| ADNI | mmyear | What is the year? |  |
| ADNI | mmmonth | What is the month? |  |
| ADNI | mmday | What day of the week is today? |  |
| ADNI | mmseason | What season is it? |  |
| ADNI | mmhospit | What is the name of this hospital (clinic, place)? |  |
| ADNI | mmfloor | What floor are we on? |  |
| ADNI | mmcity | What town or city are we in? |  |
| ADNI | mmarea | What county (district, borough, area) are we in? |  |
| ADNI | mmstate | What state are we in? |  |
| ADNI | mmball | Ball |  |
| ADNI | mmflag | Flag |  |
| ADNI | mmtree | Tree |  |
| ADNI | mmballdl | Ball delayed |  |
| ADNI | mmflagdl | Flag delayed |  |
| ADNI | mmtreedl | Tree delayed |  |
| ADNI | imm1sum | Immediate recall of the MoCA list (#1) |  |
| ADNI | imm2sum | Immediate recall of the MoCA list(#2) |  |
| ADNI | delsum | Delayed recall of the MoCA list |  |

* MoCA (blue) items were only administered in ADNI GO/2 while orange items were in all ADNI waves (1/GO/2).

* ADNI administered two versions (different word lists) of RAVLT (avtot1–avdeltot) and three different versions of ADAS-Cog items (q*) across waves. We ran the model separately for the two versions. The ADAS-Cog versions were found to be equivalent while the RAVLT versions were not. For determining secondary factor structures and extracting model fit statistics, we considered all RAVLT versions to be equivalent. The different versions of RAVLT were taken into account in the final co-calibration phase.

* There were additional MoCA items, which were the same (theoretically) as corresponding items from the Mini-Mental State Examination (MMSE). We excluded MoCA items if those items were already asked as part of the neuropsychological battery.

**ROS/MAP:** Final model was a data driven bifactor model with CFI = 0.988, TLI = 0.986, and RMSEA = 0.073. The following items were included in the CFA analysis:

| **Study** | **Variable** | **Description** | **Comments** | **Secondary Structure** |
| --- | --- | --- | --- | --- |
| ROS/MAP | Q1mme | What is the year? |  |  |
| ROS/MAP | Q2mme | What is the season of the year? |  |  |
| ROS/MAP | Q3mme | What is the date? |  |  |
| ROS/MAP | Q4mme | What is the day of the week? |  |  |
| ROS/MAP | Q5mme | What is the month? |  |  |
| ROS/MAP | Q6mme | What state are we in? |  |  |
| ROS/MAP | Q8mme | What city are we in? |  |  |
| ROS/MAP | Q7mme | What county are we in? |  |  |
| ROS/MAP | Q9mme | What room are we in? |  |  |
| ROS/MAP | Q10amme | What is the address of this place? |  | F1 |
| ROS/MAP | Q10bmme | Street Name |  | F1 |
| ROS/MAP | atb1 | Apple, table, penny (immediate) | 3 items collapsed |  |
| ROS/MAP | story | Logical memory |  | F4 |
| ROS/MAP | WordT1 | Word list learning Trial 1 | 10 items collapsed | F2 |
| ROS/MAP | WordT2 | Word list learning Trial 2 | “ | F2 |
| ROS/MAP | WordT3 | Word list learning Trial 3 | “ | F2 |
| ROS/MAP | WordRec | Which one of these words is from that list? (Word list recognition) | 10 items collapsed |  |
| ROS/MAP | Recall | Word list recall | 10 items collapsed | F2 |
| ROS/MAP | ebmt | East Boston immediate recall | 12 items collapsed | F3 |
| ROS/MAP | atb2 | apple, table, penny (delayed) | 3 items collapsed |  |
| ROS/MAP | ebdr | East Boston delayed recall | 12 items collapsed | F3 |
| ROS/MAP | Delay | Tell me the story again |  | F4 |

**Co-calibration of Memory across ACT, ADNI, ROS/MAP**

| **Study** | **Variable** | **Secondary Structure** | **Comments** |
| --- | --- | --- | --- |
| ACT, ADNI, ROS/MAP | mmyear |  | Q1mme in ROS/MAP; yr in ACT |
| ACT, ADNI, ROS/MAP | mmseason |  | Q2mme in ROS/MAP; casi_ssn in ACT |
| ACT, ADNI, ROS/MAP | mmdate | F7 | Q3mme in ROS/MAP; casi_dat in ACT |
| ACT, ADNI, ROS/MAP | mmday |  | Q4mme in ROS/MAP; day in ACT |
| ACT, ADNI, ROS/MAP | mmmonth | F7 | Q5mme in ROS/MAP; mo in ACT |
| ACT, ADNI, ROS/MAP | mmctst |  | Collapsed (Q6mme Q8mme) in ROSMAP and (mmcity mmstate) in ADNI to create a single variable; spa in ACT |
| ACT, ADNI | limmtotal | F2 | limmtotal in ADNI; w_lm_ima in ACT |
| ACT, ADNI | ldeltotal | F2 | limmtotal in ADNI; w_lm_dea in ACT |
| ROS/MAP | Q7mme |  |  |
| ROS/MAP | Q9mme |  |  |
| ROS/MAP | Q10amme | F13 |  |
| ROS/MAP | Q10bmme | F13 |  |
| ROS/MAP | atb1 |  |  |
| ROS/MAP | story | F11 |  |
| ROS/MAP | WordT1 | F10 |  |
| ROS/MAP | WordT2 | F10 |  |
| ROS/MAP | WordT3 | F10 |  |
| ROS/MAP | ebmt | F12 |  |
| ROS/MAP | WordRec |  |  |
| ROS/MAP | atb2 |  |  |
| ROS/MAP | Recall | F10 |  |
| ROS/MAP | ebdr | F12 |  |
| ROS/MAP | Delay | F11 |  |
| ACT | mat_mem | F6 |  |
| ACT | w_in_c1 | F6 |  |
| ACT | w_in_c2 | F6 |  |
| ACT | w_in_c3 | F6 |  |
| ACT | w_rcl_c | F6 |  |
| ACT | w_rcg_t | F6 |  |
| ACT | cp_re_ci |  |  |
| ACT | cp_re_di |  |  |
| ACT | cp_re_re |  |  |
| ACT | cp_re_cu |  |  |
| ACT | w_lm_imb | F3 |  |
| ACT | w_lm_deb | F3 |  |
| ACT | w_vp_ine | F4 |  |
| ACT | w_vp_inh | F5 |  |
| ACT | w_vp_ree | F4 |  |
| ACT | w_vp_reh | F5 |  |
| ACT-CASI | rgs1 |  |  |
| ACT-CASI | rc1a | F1 |  |
| ACT-CASI | rc1b | F1 |  |
| ACT-CASI | rc1c | F1 |  |
| ACT-CASI | spb |  |  |
| ACT-CASI | rc2a | F1 |  |
| ACT-CASI | rc2b | F1 |  |
| ACT-CASI | rc2c | F1 |  |
| ACT-CASI | rcobj |  |  |
| ADNI | avtot1 |  | Each RAVLT item was split into two items to account for two versions of RAVLT used in ADNI at specific waves where both versions of the same item were loaded into the same secondary structure |
| ADNI | avtot2 | F8 |  |
| ADNI | avtot3 | F8 |  |
| ADNI | avtot4 | F8 |  |
| ADNI | avtot5 | F8 |  |
| ADNI | avtot6 | F8 |  |
| ADNI | avtotb |  |  |
| ADNI | avdel30min | F8 |  |
| ADNI | avdeltot |  |  |
| ADNI | q1score | F9 |  |
| ADNI | q4score | F9 |  |
| ADNI | q7score |  |  |
| ADNI | q8score |  |  |
| ADNI | mmhospit |  |  |
| ADNI | mmfloor |  |  |
| ADNI | mmarea |  |  |
| ADNI | bft1 |  | Immediate—ball, flag, tree collapsed |
| ADNI | bft2 |  | Delayed—ball, flag, tree collapsed |
| ADNI | imm1sum |  |  |
| ADNI | Imm2sum |  |  |
| ADNI | delsum |  |  |

**Addition of A4:** As detailed above, items where the A4 item was the same as an item with parameters from the ACT/ADNI/ROS-MAP analyses, we used those previously co-calibrated parameters. For items administered only to A4 participants, we freely estimated item parameters from the data set.

| **Study** | **Variable** | **Description** | **Secondary Structure** |
| --- | --- | --- | --- |
| ACT, ADNI, ROS/MAP, A4 | mmyear | What is the year? |  |
| ACT, ADNI, ROS/MAP, A4 | mmseason | What season is it? |  |
| ACT, ADNI, ROS/MAP, A4 | mmdate | What is today’s date? | F1 |
| ACT, ADNI, ROS/MAP, A4 | mmday | What day of the week is today? |  |
| ACT, ADNI, ROS/MAP,A4 | mmmonth | What month is it? | F1 |
| ACT, ADNI, ROS/MAP, A4 | mmctst | Collapsed: What city are we in? What state are we in? |  |
| A4 | fctott1 | FCSRT* Trial 1 total | F2 |
| A4 | fctott2 | FCSRT* Trial 2 total | F2 |
| A4 | fctott3 | FCSRT* Trial 3 total | F2 |
| A4 | limmtot_rm | Logical Memory (Robert Miller Story) | F3 |
| A4 | ldeltot_rm | Logical Memory (Robert Miller Story) | F3 |
| ACT, ADNI, A4 | limmtotal | Logical Memory (Anna Thompson Story) | F4 |
| ACT, ADNI, A4 | ldeltotal | Logical Memory (Anna Thompson Story) | F4 |
| ADNI, A4 | mmhospit | What is the name of this hospital (clinic, place)? |  |
| ADNI, A4 | mmfloor | What floor are we on? |  |
| ADNI, A4 | mmarea | What county (district, borough, area) are we in? |  |
| ADNI, A4 | bft1 | Immediate—ball, flag, tree collapsed |  |
| ADNI, A4 | bft2 | Delayed—ball, flag, tree collapsed |  |

*FCSRT = Free and Cued Selective Reminding Test

#### 3B. Executive Function

**ACT:** Final model was a data driven bifactor model with CFI = 0.948, TLI = 0.929, and RMSEA = 0.064. The following items were included in the CFA analysis:

| **Study** | **Variable** | **Description** | **Comments** | **Secondary Structure** |
| --- | --- | --- | --- | --- |
| ACT | mat_attn | Mattis Dementia Rating Scale, Attention score |  |  |
| ACT | mat_conc | Mattis Dementia Rating Scale, Concentration score |  |  |
| ACT | mat_ip | Mattis Dementia Rating Scale, initiation / perseveration score |  |  |
| ACT | tr_a_tm | Trails A |  | F1 |
| ACT | tr_b_tm | Trails B |  | F1 |
| ACT | clockdr | Clock |  |  |
| ACT-CASI | dbsum | repeat numbers backward 1–3 | Repeat numbers backward—3 trials collapsed | F2 |
| ACT-CASI | subtra | Subtraction 1–3 | Subtraction—3 trials collapsed | F2 |
| ACT-CASI | sim | similarities |  |  |
| ACT-CASI | jgmt | judgement |  |  |

**ADNI:** Final model was a theory driven methods-effects bifactor model with CFI = 0.951, TLI = 0.946, and RMSEA = 0.041. The following items were included in the CFA analysis:

| **Study** | **Variable** | **Description** | **Comments** | **Secondary Structure** |
| --- | --- | --- | --- | --- |
| ADNI | clockcirc | Approximately circular face |  |  |
| ADNI | clocksym | Symmetry of number placement |  | F2 |
| ADNI | clocknum | Correctness of numbers |  | F2 |
| ADNI | clockhand | Presence of the two hands |  |  |
| ADNI | clocktime | Presence of the two hands, set to ten after eleven |  |  |
| ADNI | dspanbac | Backward Total Correct |  | F4 |
| ADNI | traascor | Part A Time to Complete |  | F3 |
| ADNI | trabscor | Part B Time to complete |  | F3 |
| ADNI | digitscor | Digit Symbol Total Correct |  | F1 |
| ADNI | dspanfor | Digit Span Forward Total Correct |  | F4 |
| ADNI | q13score | Number cancellation task |  | F1 |
| ADNI | absmeas | Abstraction: watch-ruler |  |  |
| ADNI | abstran | Abstraction: train-bicycle |  |  |
| ADNI | trails | MoCA Trails |  |  |
| ADNI | digback | Digits Backward | 5 trials collapsed |  |
| ADNI | serial | Serial 7 total |  |  |
| ADNI | digfor | Digits Forward |  |  |
| ADNI | letters | List of Letters/Tapping: # Errors |  |  |

**ROS/MAP:** Final model was a theory driven methods-effects model with CFI = 0.975, TLI = 0.960, and RMSEA = 0.064. The following items were included in the CFA analysis:

| **Study** | **Variable** | **Description** | **Comments** | **Secondary structure** |
| --- | --- | --- | --- | --- |
| ROS/MAP | AA | Which piece would complete the pattern… | 4 A patterns merged |  |
| ROS/MAP | BB | Which piece would complete the pattern… | 8 B patterns merged |  |
| ROS/MAP | Q12bmme | Spell WORLD backwards |  |  |
| ROS/MAP | DigBak | digits backward | combined 12 items |  |
| ROS/MAP | cts_sdmt | symbol digits modality (oral) |  | F1 |
| ROS/MAP | cts_nccrtd | Number comparison |  | F1 |
| ROS/MAP | DigFor | digits forward | combined 12 items |  |

**Co-calibration of executive functioning across ACT, ADNI, ROS/MAP:**

| **Study** | **Variable** | **Description** | **Secondary structure** |
| --- | --- | --- | --- |
| ACT, ADNI | traascor | Trails A | F3 |
| ACT, ADNI | trabscor | Trails B | F3 |
| ADNI, ROS/MAP | dspanfor | Digit Span Forward: Total Correct | F4 |
| ADNI, ROS/MAP | dspanbac | Digit Span Backward: Total Correct | F4 |
| ROS/MAP | AA | Which piece would complete the pattern… |  |
| ROS/MAP | BB | Which piece would complete the pattern… |  |
| ROS/MAP | Q12bmme | Spell WORLD backwards |  |
| ROS/MAP | cts_sdmt | symbol digits modality (oral) | F6 |
| ROS/MAP | cts_nccrtd | Number comparison | F6 |
| ACT | mat_attn | Mattis Dementia Rating Scale |  |
| ACT | mat_conc | Mattis Dementia Rating Scale |  |
| ACT | mat_ip | Mattis Dementia Rating Scale |  |
| ACT | clockdr | Clock |  |
| ACT-CASI | dbsum | repeat numbers backward | F5 |
| ACT-CASI | subtra | subtraction | F5 |
| ACT-CASI | sim | similar types |  |
| ACT-CASI | jgmt | judgement |  |
| ADNI | clockcirc | Approximately circular face |  |
| ADNI | clocksym | Symmetry of number placement | F2 |
| ADNI | clocknum | Correctness of numbers | F2 |
| ADNI | clockhand | Presence of the two hands |  |
| ADNI | clocktime | Presence of the two hands, set to 10 after 11 |  |
| ADNI | digitscor | Digit Symbol Total Correct | F1 |
| ADNI | q13score | Number cancellation task | F1 |
| ADNI | absmeas | Abstraction: watch–ruler |  |
| ADNI | abstran | Abstraction: train–bicycle |  |
| ADNI | trails | Trails |  |
| ADNI | digback | Digits Backward |  |
| ADNI | serial | Serial 7 |  |
| ADNI | digfor | Digits Forward |  |
| ADNI | letters | List of Letters/Tapping: # Errors |  |

**A4:** Sufficient data was not available in A4 to create a co-calibrated executive function score.

### Supplemental Text References

1	Hu Lt, Bentler PM (1999) Cutoff criteria for fit indexes in covariance structure analysis: Conventional criteria versus new alternatives. Structural Equation Modeling: A Multidisciplinary Journal 6: 1-55 Doi 10.1080/10705519909540118

2	Muthén LK, Muthén BO (1998-2015) Mplus User's Guide. Seventh Edition. Muthén & Muthén, Los Angeles, CA.

3	Reeve BB, Hays RD, Bjorner JB, Cook KF, Crane PK, Teresi JA et al (2007) Psychometric evaluation and calibration of health-related quality of life item banks: plans for the Patient-Reported Outcomes Measurement Information System (PROMIS). Med Care 45: S22-31 Doi 10.1097/01.mlr.0000250483.85507.04
